# A biomechanical model for the relation between bite force and mandibular opening angle in arthropods

**DOI:** 10.1101/2022.08.17.504316

**Authors:** Frederik Püffel, Richard Johnston, David Labonte

## Abstract

Bite forces play a key role in animal ecology: they affect mating behaviour, fighting success, and the ability to feed. Although feeding habits of arthropods have an enormous ecological and economical impact, we lack fundamental knowledge on how the morphology and physiology of their bite apparatus controls bite performance and its variation with mandible gape. To address this gap, we derived a comprehensive biomechanical model that characterises the relationship between bite force and mandibular opening angle from first principles. We validate the model by comparing its geometric predictions with morphological measurements on CT-scans of *Atta cephalotes* leaf-cutter ants. We then demonstrate its deductive and inductive power with three exemplary use cases: First, we extract the physiological properties of the leaf-cutter ant mandible closer muscle from *in-vivo* bite force measurements. Second, we show that leaf-cutter ants are extremely specialised for biting: they generate maximum bite forces equivalent to about 2600 times their body weight. Third, we discuss the relative importance of morphology and physiology in determining the magnitude and variation of bite force. We hope that our work will facilitate future comparative studies on the insect bite apparatus, and advance our knowledge of the behaviour, ecology and evolution of arthropods.

## Introduction

Bite forces are a performance metric paramount to animal behaviour, ecology and evolution [1]. They determine access to food sources [2–7], fighting ability, reproductive success [8–11], and the ability to escape predators [12]. Strikingly, the vast majority of bite force studies has been conducted on vertebrates and crustaceans [e. g. 2–8, 11, 13–52]. In sharp contrast, comparatively few studies exist for the hyperdiverse non-crustacean arthropods [but see 9, 10, 12, 53–63].

This disparity is as striking as it is surprising; arthropods rely on bite forces just as much as vertebrates: to fight over access to food and mating partners [9, 10], and in other key behaviours such as nest building [64]. The arguably most important role of bite forces in arthropods, however, is that they determine the ability to mechanically process food [65–67]. Indeed, plant-eating insects have an enormous impact on our lives – they destroy around 11 % of our crops, causing billions of US dollars in economic damage, and may jeopardise global food security in a warming climate [68–72] – and affect entire ecosystems [73], To provide but two illustrative examples, leaf-cutter ants accelerate the cycling of nutrients in the Neotropics through massive defoliation and decomposition of plant material [74], and seed-harvesting ants increase the dispersion rate and regeneration of myrmecochorous plants [75]. The need to mechanically process plant-material to feed is likely one of the driving factors in the evolutionary arms race between plants and insects, and thus constitutes a significant aspect of insect diversity [76–78].

Despite this broad significance, fundamental knowledge about bite performance across the arthropod tree of life is scarce. A reasonable and indeed common starting point to characterise bite performance within and between species may be the magnitude of the peak bite force [79]. This peak force, however, can be difficult to measure, because arthropods are often small; it is also challenging to predict, as it depends on the geometry of the bite apparatus and the physiology of muscle, and both vary substantially across arthropods [23, 78].

To make matters worse, bite force typically varies with the mandibular opening angle or gape, adding further complexity [4, 7, 10, 26, 38, 44, 48, 56, 60, 80]. This variation is not merely a complication, but is functionally relevant and may reflect species-specific demands. For example, masticating bats produce largest bite forces at small opening angles [26, 48], whereas snail-eating carps produce maximum forces at comparatively larger angles [4]; predatory king salmons produce maximum bite force at a relatively larger mandible gape than filterfeeding pink salmons [7]; and stag beetles generate largest bite forces at opening angles that are typical during combat [10]. The ecological and behavioural needs of each species hence likely drive the morphological and physiological properties of the musculoskeletal bite apparatus which determine the variation of bite force with mandibular opening angle.

Although bite forces depend on a large number of anatomical and physiological parameters, they are of mechanical origin, which makes them amenable to exact analysis from first principles. A quantitative model which predicts the magnitude and variation of bite force with opening angle from morphology and physiology of the bite apparatus would provide a powerful tool to study comparative bite performance and head anatomy, and may thus increase our understanding of arthropod behaviour, ecology and evolution. In the following paragraphs, we derive such a model and then validate it using *Atta cephalotes* leafcutter ants as model organism. We demonstrate the utility of the model by extracting the force-length properties of the mandible closer muscle from *in-vivo* bite force measurements, discuss the morphology and exceptional performance of the leaf-cutter ant bite apparatus in an ecological and comparative context, and discuss the possibility and accuracy of ‘minimal’ bite force models which allow colleagues to predict the magnitude of the bite force and its variation with opening angle from a reduced set of accessible parameters.

### A biomechanical analysis of biting in arthropods

The magnitude of the bite force, |**F**_*b*_|, is determined by the architecture and the physiological properties of the musculoskeletal bite apparatus. These two components are fully captured by two simple terms: the net muscle force which pulls on the apodeme (from here on apodeme force); and the mechanical advantage of the force-transmission system, *MA*, i. e. the ratio of two characteristic moment arms [53, 56, 81]. The net muscle force is the vector sum of the forces generated by individual fibres. Combined, all fibres produce a force with a magnitude equal to the product between a characteristic muscle stress, *σ*, and a characteristic cross-sectional area, *A_phys_*. However, unless the fibres are perfectly aligned, only some fraction of the fibre force magnitude contributes to the magnitude of the net force. This fraction is typically characterised by an average angle of pennation, *φ* (throughout this manuscript, all vectors are bold, and all unit vectors are indicated by a hat). We may thus write:

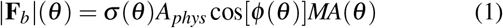

We note that only one of the four key parameters in this equation, *A_phys_*, is independent of the mandibular opening angle *θ*, defined here as the angle between the lateral head axis and the projection of the outlever on the plane of rotation [see Fig. 1A-C and 82]. In the following paragraphs, we derive the critical functions *σ*(*θ*), cos[*φ*(*θ*)] and *MA*(*θ*) from first principles. Understanding how the morphology and physiology of the bite force apparatus control the variation of bite force with opening angle is not just an enjoyable exercise in mechanics, but has substantial biological implications, for it determines the accessibility of food items of different size [e. g. 4], and potentially prey handling times [44]. However, this problem has received relatively little quantitative attention [e. g. 4, 10, 44, 80, 82].

**Figure 1.**
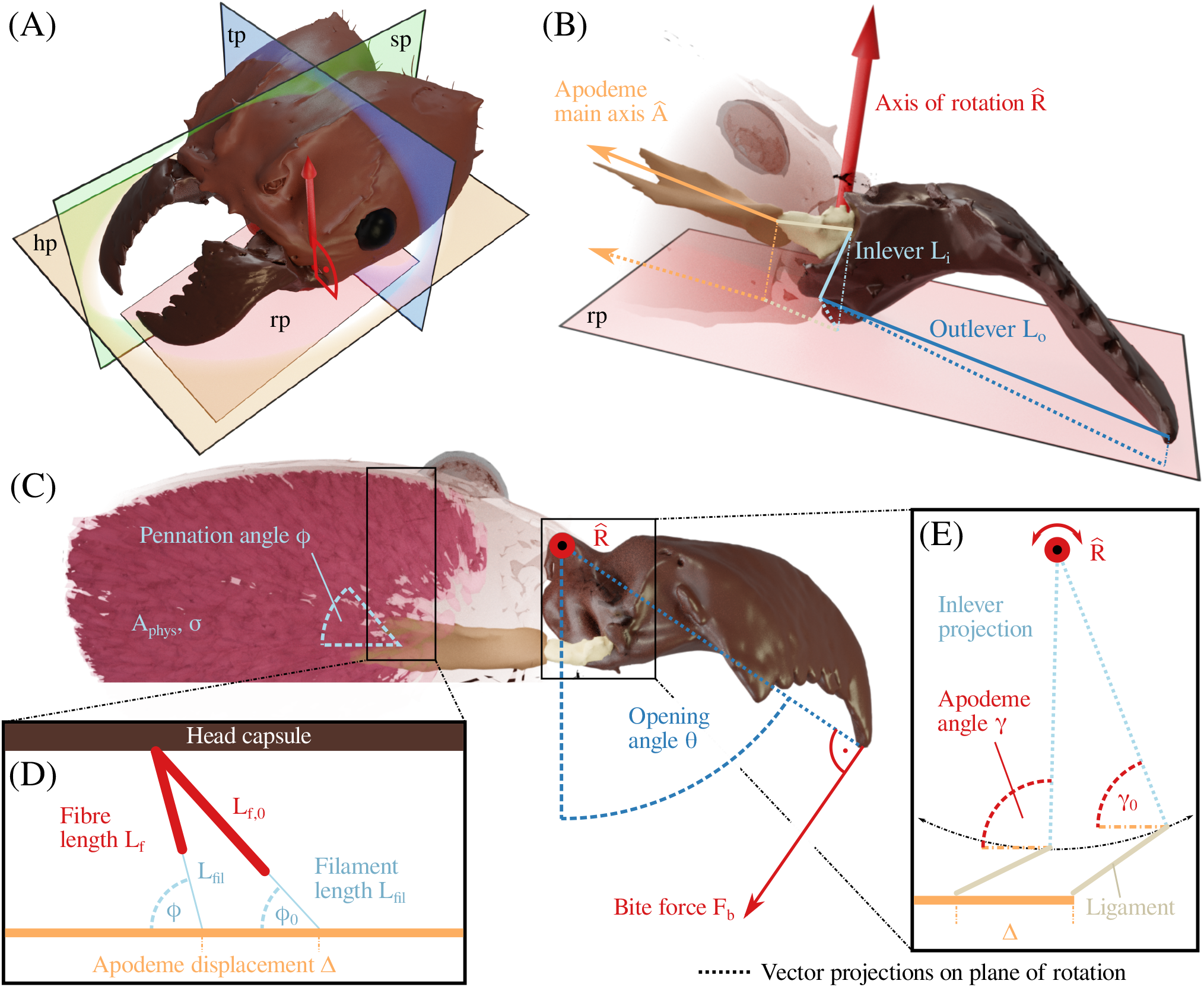
(**A-C**) The bite force **F**_b_ is a function of muscle stress *σ*, physiological cross-sectional area *A_phys_*, pennation angle *φ*, apodeme angle *γ*, and the mechanical advantage determined by inlever **L**_*i*_ and outlever **L**_*o*_, the axis of rotation 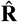, and the apodeme main axis 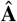. The mechanical advantage and muscle stress, as well as the apodeme and pennation angle, vary with the mandibular opening angle *θ*, so affecting the magnitude of the maximum bite force that can be generated at different opening angles. *γ* and *θ* are defined with respect to the vector projections of 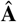, **L**_*i*_ and **L**_*o*_ onto the rotational plane (rp). The position of this plane with respect to the head coordinate system, defined by the horizontal (hp), transversal (tp) and sagittal (sp) plane, can often be inferred from joint morphology [see 81]. (**D**) Muscle fibres attach either directly or via thin filaments to the apodeme. When muscle fibres shorten (*L*_*f*,0_ to *L_f_*), the pennation angles increase (*φ*_0_ to *φ*), and the apodeme is displaced (**Δ**); the filament length remains approximately constant (see text). (**E**) As a result of the apodeme displacement, the inlever rotates about 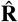 and *γ* changes. Notably, the apodeme appears to move in pure translation, i. e. its main axis has a constant orientation. The lateral displacement of the attachment point which must accompany mandible rotation is likely accommodated by rotation of the putatively soft apodeme ligament, which connects the apodeme to the mandible base.

We derive bite force as a function of the opening angle because this relationship is biologically meaningful and directly relates to the typical experimental procedure to measure bite forces [e.g. 22, 26, 60]. Physically, however, the bite force is not a function of the opening angle, but of the muscle activation and the muscle fibre stretch ratio *λ*. The causal relationship |**F_*b*_**| = *f* (*λ*) is provided in the SI. Throughout the derivation, we use the following assumptions: mandible kinematics are fully characterised by a single axis of rotation; the tendon-like apodeme connects to the mandible base via a single attachment point and displaces along its main morphological axis which coincides with the net muscle force vector; and the head capsule, apodeme and mandible are rigid. Due to the abundance of dicondylic mandible joints across the insecta, which are putative hinge joints [83, 84], and the comparatively high elastic modulus of sclerotised cuticle [Young’s modulus ≥ 1 GPa, 85, 86], these assumptions likely hold for the vast majority of all insect species with chewing-biting mouthparts, as well as many other arthropods. The model thus has considerable generality, but we stress that it may not be readily applicable to vertebrate biting, which may involve multiple muscle insertion points, non-rigid tissue deformation, jaw articulations with multiple degrees of freedom, and more complex muscle deformations [87].

#### Mechanical advantage

Musculoskeletal lever systems are often characterised in terms of their mechanical advantage; the ratio between two characteristic moment arms [e.g. see 10, 53, 67]. These moment arms are determined by the location of two force transmission points, the joint centre, and the orientation of the relevant force vectors and the joint rotational axis (see Fig. 1).

The points of force transmission are readily identified: *internally*, it is the attachment point of the apodeme to the mandible base (in most insects, muscle force is transmitted to the mandible via a single closer apodeme [84]); *externally*, it is the bite contact point on the mandibular blade. The rotational axis, 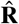, is determined by joint morphology, and may often be inferred from the axis connecting the two joint condyles [10, 61, 62, 82, 88]. We use the apodeme main axis 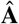 as proxy for the orientation of the *internal* apodeme force vector, because it is easier to measure and is likely closely aligned with the net muscle force vector [81]; where apodemes have multiple well-developed “branches”, a more suitable proxy may be derived from muscle geometry instead [see e.g. 10, 82]. The *external* bite force vector **F**_*b*_, in turn, is perpendicular to both the rotational axis and the mandible outlever by definition (see Fig. 1C).

To calculate the mechanical advantage, we extract the shortest distance between 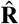 and the points of force transmission. To this end, we first define a specific plane of rotation by choosing a point on the rotational axis as the joint centre, in order to then find the planar (2D) components of the relevant 3D position vectors (see Fig. 1A). In theory, the choice of this point is arbitrary as it does not affect the moment arm calculation. In practice, however, it can be informed by joint morphology and may, in fact, be used to define the physical location of 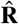, e. g. by using the centre of a joint condyle. Second, we define the in-and outlever, **L**_*i*_ and **L**_*o*_, respectively, as the vectors connecting the joint centre with the internal and external points of force transmission. We project these vectors onto the plane of rotation and calculate the length of the projections, 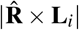 and 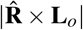, respectively (see Fig. 1B); these lengths are equal to the shortest distance between the axis of rotation and the two points of force transmission.

The length of the projected in- and outlever represents the maximum possible moment arm. The effective lever length, however, may be shorter, depending on its orientation relative to the relevant force vector. Because the bite force is perpendicular to the outlever by definition, **F**_*b*_ rotates with **L**_*o*_ as the mandible opens and closes; it follows that 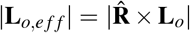 for all opening angles.

For the inlever, in contrast, the geometric dependency is considerable more complex. The orientation of the apodeme force remains approximately constant across mandibular opening angles, because the apodeme does not rotate. However, because the joint converts apodeme translation into mandible rotation, the angle between inlever and 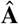 changes with opening angle (see Fig. 1C,E). Calculating the magnitude of the effective inlever |**L**_*i,eff*_| at different opening angles thus requires to define the angle *γ* between the vector projections of inlever and apodeme main axis onto the plane of rotation. *γ* can be written as a direct function of opening angle, *γ*(*θ*) = *θ*_0_ - *θ* + *γ*_0_, where *γ*_0_ is measured at an arbitrary reference opening angle *θ*_0_ [for a similar calculation, see 10, 80, 82]. For *γ* ≈ 90°, the effective inlever takes a maximum value of 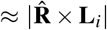, and a small change in opening angle will be associated with a small change in effective inlever. If *γ* is acute or obtuse, however, the effective inlever is small, and rapidly varies with opening angle. These results may be understood in analogy to the effect of pennation angle on apodeme displacement (see below): both angles control the amount of rotation associated with a unit of translational displacement. The two characteristic angles *φ* and *γ* thus crucially determine the gearing of the muscle force across opening angles. However, the importance of *γ* in this context has received comparatively little attention [but see 60].

As a last step in the calculation of the mechanical advantage, we account for the fact that the apodeme force vector may not lie in the plane of rotation. The orientation of 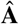 relative to 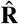 matters because only the force components in the plane of rotation can be transmitted through the mandible joint; all other components result in joint reaction forces instead. We calculate the fraction of the force acting in the plane of rotation as the cross-product between the two relevant unit vectors, 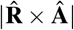. The magnitude of the effective inlever as a function of the opening angle then follows as:

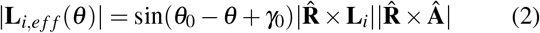

We define the mechanical advantage as the ratio between effective in- and outlever, *MA* = |**L**_*i,eff*_|/|**L**_*o,eff*_|.

#### Apodeme displacement

Any rotation of the mandible is associated with a displacement of the apodeme attachment point. This displacement has two components in the plane of rotation: one longitudinal component along the projection of the apodeme main axis, and one lateral component perpendicular to it. Previous work has noted that the lateral component is small for small changes in opening angle, and can thus be neglected [82]. This simplifying assumption, based on the small angle approximation, holds if the length of the inlever projection onto the plane of rotation is small, and the apodeme angle remains close to 90° throughout the opening range. However, for large variations in opening angle, it is inaccurate. As an illustrative example, if *θ* changes by 40°, the lateral displacement is about one-third of the longitudinal displacement, 1/3 ≈ (1 - sin50°)/cos50° (see Fig 1E). The longitudinal displacement is likely associated with an equivalent displacement of the apodeme along its main axis; the lateral displacement, however, is likely not: If the apodeme was to translate laterally or rotate significantly, a fraction of the closer muscle would lengthen considerably while another fraction may be slack. Indeed, the apodeme would eventually hit the head capsule. Lateral translation or rotation of the apodeme is thus biologically implausible, and to the best of our knowledge, has never been reported. However, lateral displacements of the attachment point must occur, because it moves along a circular path (see Fig. 1E).

We hypothesise that these lateral displacements are enabled by the rotation of the apodeme ligament, a putatively flexible connective tissue which bridges the sclerotised apodeme and the mandible [89–91, see Fig. 1E]. It remains unclear how exactly the apodeme, ligament, and mandible are mechanically linked. We conjecture that both longitudinal *and* transversal forces can be transmitted from the apodeme through the ligament, so effectively shifting the apodeme force vector to the point where the ligament attaches to the mandible. At this attachment point, we assume a hinge-like connection, which enables variation in *γ*.

Based on the above hypothesis, the displacement of the apodeme along its main axis, Δ, is equal to the longitudinal displacement of the apodeme attachment point corrected by the misalignment between 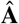 and the plane of rotation; in analogy to the effective inlever, only apodeme displacement components in the plane of rotation contribute to mandible rotation. The associated out-of-plane displacements are likely also enabled by the ligament (and may typically be very small, see below). As a consequence and in analogy to Eq. 2, the apodeme displacement follows as:

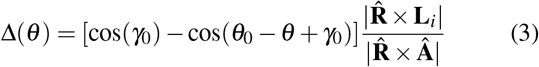

We define Δ such that it increases as the mandible closes; it can however be defined with respect to any reference opening angle *θ*_0_, for Δ(*θ*_0_) = 0. Similar to the effective inlever, the change of Δ with *θ* depends on the apodeme angle *γ*. If *γ* ≈ 90°, a small change in opening angle will be associated with a large apodeme displacement; if *γ* is acute or obtuse, the opposite holds.

#### Pennation angle

All muscle fibres attach to the rigid head capsule on one side, and insert onto the surface of the apodeme on the other. Consequently, muscle fibres that attach at a reference angle φ_0_ > 0° with respect to the apodeme main axis change their relative orientation when they shorten to move the apodeme – they rotate. The associated change in pennation angle φ, first derived by [92], depends on φ_0_, and the average distance between muscle fibre origins and insertion points, *L*_*t*,0_ (see Fig. 1D and SI for derivation):

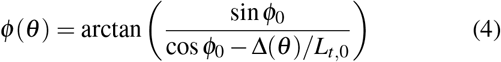

#### Muscle stress

Muscle stress is a function of fibre length, as the relative overlap and lattice spacing between thick and thin myofilaments varies when fibres are stretched or contract [93–95]. In contrast to all other parts of the model, this force-length relationship cannot be easily derived from first principles [but see 96]. Instead, it has been modelled with a variety of empirical functions; one simple and thus attractive choice is a Gaussian probability density function [e. g. 97, 98]:

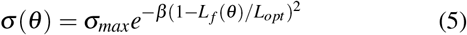

This function has a maximum *σ_max_* at an optimal fibre length *L_f_* = *L_opt_*; *β* is a shape parameter which determines how quickly the stress decays as the fibre is stretched or contracts. Eq. 5 contains only three parameters that cannot be easily measured from CT scans: *σ_max_*, *L_opt_* and *β* – they are physiological parameters which vary with the microanatomy of the muscle, such as the relative length of the myofilaments [94, 99]. Note that *σ_max_*, *L_opt_* and *β* may vary within the muscle between directly- and filament-attached fibres [100, and see below], but the form of Eq. 5 holds for both. We stress that more complex functions of *σ*(*θ*), e. g. those which include passive muscle forces [97, 101], may characterise the variation of muscle stress with opening angle more accurately, but at the cost of a larger number of parameters that need to be quantified (see below).

#### Fibre length

To predict muscle stress, the relationship between muscle fibre length and opening angle needs to be known. In insects, this relationship has previously been modelled with a simple linear function [10], which may be sufficiently accurate provided that the range of opening angles is small. For a large range of opening angles, however, the small angle approximation is inaccurate, and the functional relation is more complex. Consider a muscle fibre of total length *L_t_* which attaches directly to the apodeme. The variation of fibre length *L_d_* with apodeme displacement is determined by the reference fibre length *L*_*d*,0_, the fibre pennation angle φ_0_, and Δ(*θ*) [see 102, and SI for derivation]:

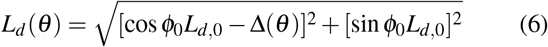

However, in some insects, fibres may attach via thin cuticular filaments instead [81, 103]. Filament-attachment increases volume occupancy and cross-sectional area of the muscle while keeping the apodeme compact; it is a light-weight solution to strongly increase bite force capacity in the limited space provided by the rigid head capsule [81, 104]. In addition to these *static* effects, filament-attachment also has *dynamic* effects: Consider two muscle fibres of equal length; one is directly- and one filament-attached. The same fibre shortening will lead to a different apodeme displacement, because filaments reduce the amount of rotation per unit fibre strain (see Fig. 1D). As a consequence, the change in pennation angle associated with fibre-shortening is smaller in the filament-attached fibre.

In analogy to Eq. 6, the relation between the length of filament-attached fibres and apodeme displacement depends on the initial fibre length *L*_*f*,0_, pennation angle *φ*_0_, Δ(*θ*) and filament length *L_fil_* as (see SI for derivation):

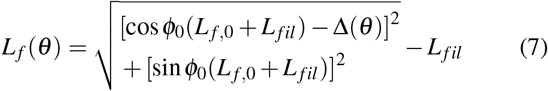

where we assumed that filament length is independent of pennation angle (the strain in the filaments across the opening range is less than 2%, see SI).

The last parameter in Eq. 1 is the physiological crosssectional area of the muscle, *A_phys_*. The physical cross-sectional area of the muscle changes as fibres change length because muscle is approximately incompressible. *A_phys_*, however, is a characteristic cross-sectional area, and thus invariant to strain. In vertebrates, *A_phys_* is often measured for a relaxed muscle [e. g. 105], but this definition is problematic in comparative work, because the relaxed fibre length may be a different fraction of *L_opt_* in different muscles. In order to avoid a systematic under- or overestimation of *σ_max_*, we thus suggest to define *A_phys_* as the physiological cross-sectional area at an equivalent point of the force-length curve, and the natural choice for this point is the cross-sectional area for fibres of length *L_opt_* at which stress is maximal.

Based on the derived geometric relations, we substitute the place-holding functions of *θ* in Eq. 1, *MA*(*θ*), *φ*(*θ*), and *σ*(*θ*), with Eq. 2 (divided by |**L**_*o,eff*_|), Eq. 4, and Eq. 5, respectively. Muscle stress is expressed as a function of fibre length, so that Eq. 6 (for directly-attached fibres) and Eq. 7 (for filament-attached fibres) need to be inserted into Eq. 5. Total muscle stress is equal to the sum of the fractional contributions of directly- and filament-attached fibres. Last, fibre length and pennation angle are expressed as functions of the apodeme displacement, so that Eq. 3 needs to be inserted into Eq. 4, Eq. 6 and Eq. 7. The resulting equation contains a total of 20 parameters, including *θ*.

The above model fully captures the determinants of bite force in arthropods and its variation with opening angle. Practically, the bite force is rarely measured directly. Instead, its magnitude, |**F**_*b*_|, is inferred from measurements with 1D or 2D force sensors [e. g. see 22, 53, 60, 106]. As a consequence, at least one of the vector components has to be inferred from geometry. For 1D sensors, the key geometric variable is the angle *α* between the force sensitive axis, and the bite force, **F**_*b*_; the measured force, **F**_*m*_, is equal to the projection of **F**_*b*_ onto the sensitive axis:

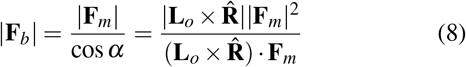

where the term 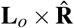 represents the orientation of **F**_*b*_. The inner product between 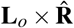 and **F_*m*_**, divided by their respective magnitudes, is equal to cos *α*. The sensitive axis is determined by the sensor design; the orientation of **F**_*b*_, however, is subject to experimental variation, for it depends on the orientation of the mandible outlever relative to the axis of rotation (see above). The corresponding relationship between |**F**_*b*_| and |**F_*m*_**| for 2D sensors is provided in the SI.

## Materials & methods

### Study animals

In order to validate the derived geometric relations (Eqs. 2–7), we extracted the morphological determinants of bite force from insect heads of similar size, but with a wide variation in mandibular opening angles. We then used the validated geometric model to extract the force-length properties of mandible closer muscles by conducting *in-vivo* bite force measurements. For both sets of experiments, we use leaf-cutter ants (genus *Atta*). Leaf-cutter ants constitute an excellent model organism for at least three reasons. First, biting is key to their ecology; they cut food sources of varying thickness and toughness to feed a fungus grown as crop [107, 108]. Second, the mandible closer muscle consists of both directly- and filament-attached muscle fibres [103], which enables us to verify the two separate geometric relations between fibre length and apodeme displacement (Eqs. 6 and 7). Third, mandibular opening angles observed during natural behaviour span a large range of around 70^°^ (see below), allowing us to validate the model across a maximum opening angle range.

Ants were collected from an *Atta cephalotes* colony, kept in a climate chamber (FitoClima 12.000 PH, Aralab, Rio de Mouro, Portugal) at 25 °C and 60% relative humidity, with a 12/12 h light-dark cycle. The colony was fed with bramble, laurel, cornflakes and honey water *ad libitum*. We selected only majors with a body mass between 50 and 60 mg in order to minimise variation due to size, reduce the complexity of handling small individuals during the experiments, and to enable bite force measurements across a maximal opening range. All individuals were weighed to the nearest 0.1 mg after collection (Explorer Analytical EX124, max. 120 g x 0.1 mg, OHAUS Corp., Parsippany, NJ, USA).

### Tomography and morphometric analysis for model validation

#### Sample preparation and scanning

To obtain information on the morphological determinants of bite force, we conducted tomographic scans of five ant heads (body mass 55.0±3.8 mg), prepared such that the mandibular opening angles approximately spanned the naturally observed range. In order to experimentally control the opening angle, live ants were clamped using a 3D printed device, and offered PLA rods of varying thickness to bite onto. The rods were positioned asymmetrically between the mandibles, so that the opening angles between left and right mandible differed (see SI for details). This procedure was followed for four of the five ants; one ant was prepared without PLA rod, which resulted in mandibles that were maximally closed. We thus collected data for ten opening angles across the entire opening range. Ants were then sacrificed by freezing, causing muscles to forcefully contract, decapitated using micro scissors, and fixed in paraformaldehyde solution (4% in PBS, Thermo Fisher Scientific, Waltham, MA, USA). After 90-95h, the samples were transferred to 100% ethanol via a series of dehydration steps in 70, 80 and 90 % ethanol solutions for an hour each. To prepare the heads for CT scanning, they were stained with 1 % iodine in ethanol for at least 4.5 days [see 109]. The samples were analysed via X-ray microscopy using a Zeiss Xradia Versa 520 X-ray microscope (Carl Zeiss XRM, Pleasanton, CA, USA; for more details, see SI).

The tomographic image stacks were reoriented such that the lateral, dorso-ventral and anterior-posterior head axes aligned with the coordinate system internal to Fiji [110, for more details, see 81]. Tissue segmentation of head capsule, mandible closer muscle and closer apodemes were performed in ITK-SNAP [v3.6, 111].

#### Morphometry

From the segmented tomographic scans, a series of morphological parameters relevant to the generation of bite forces were extracted: mechanical advantage, mandibular opening angle and the corresponding mandibular gape, the rotational axis of the mandible joint, apodeme orientation and displacement, volume of the mandible closer muscle, as well as average muscle fibre length and average pennation angle.

##### Mechanical advantage

The mechanical advantage was calculated as defined by Eq. 2. The relevant lengths, **L**_*i*_ and **L**_*o*_, and angles, *γ*_0_ and *θ*0, were extracted separately for each head hemisphere. The joint centre was placed at the ventral articulation of the joint [for an exact definition, see 81]. The outlever may be defined with respect to any point on the mandibular cutting edge; we used the most distal tooth tip to provide an upper bound, **L**_*o*_ = **L**_*o,d*_. We calculated the mandible gape as twice the shortest distance between the distal tooth tip and the sagittal plane. The gape is thus negative when mandibles overlap.

##### Axis of rotation

In order to obtain the orientation of the rotational joint axis, 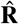, we invoke a simple result from rigid body kinematics: If the mandible rotates about a single axis, then the angle between any vector which connects two points on the mandible and 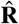 remains constant throughout mandible motion. 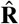 was thus determined by minimising the squared angular residuals between 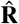 and three such vectors, **L**_*i*_, **L**_*o,d*_, and the outlever to the most proximal tooth tip, **L**_*o,p*_, using a numerical solver in python [Kang, Püffel and Labonte, in preparation]. In order to integrate vector coordinates from different scans into the same coordinate system, the image origins were first transposed to the respective joint centres. Second, the coordinates were normalised with head length, defined as anterior-posterior distance between mandible joint and the rear of the head capsule. Last, to combine data from the left and right head hemisphere, the coordinates of the latter were mirrored across the sagittal plane.

##### Apodeme displacement

In order to obtain the apodeme displacement, we first extracted the apodeme centres-of-masses via 3D particle analysis in Fiji [for more details, see 81]. The centre-of-mass coordinates were transposed and normalised as described for the rotational axis to integrate results from different individuals into the same coordinate system. Next, we performed a principal component analysis on the normalised coordinates; the first component explained the dominant part of the variation of coordinates, *R*^2^ = 0.931, and was thus selected as the axis of displacement. We defined the apodeme displacement as the distance between the normalised coordinates along this axis, multiplied with the average head length. In support of the model assumption, the axis of displacement is close to the apodeme main axis 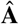, defined as in [81]; it differs only by 8 ± 2° (mean±standard deviation), so that the net force is effectively aligned with 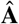 (cos(8°) ≈ 1). In the following, both axes are hence treated as equivalent.

##### Muscle architecture

The muscle volume was directly measured via ITK-SNAP. We have previously described an automated algorithm to extract fibre length and pennation angles from segmented tomography scans [81]. However, this tracing algorithm was developed for scans with a resolution and contrast only achievable with synchrotron scanning, to which we did not have access for this work. In addition, preliminary inspection of the scans revealed little ‘free volume’ between fibres, on which the automated tracing algorithm relies. We thus developed an alternative method, based on a simple sorting algorithm. This algorithm leverages the observation that all muscle fibres originate from the internal surface of the head capsule, and connect as approximately straight lines to a central apodeme either directly, or via thin sclerotised filaments [81, 103]. Obtaining pennation angle and fibre length with this algorithm involves three simple steps (see SI).

First, the 3D location of the fibre attachment points (seeds) on the head capsule are identified using Fiji as described previously [see 81, for details]. Second, each fibre seed is connected to its insertion point on the apodeme surface. To identify the insertion point, we assume that the density of fibre attachments on the apodeme is approximately constant along its length, and that muscle fibres rarely cross. On the basis of these assumptions, the fibre insertion point can be found by sorting fibre seeds and apodeme surface points by their anterior-posterior positions, to then match fibre seeds and apodeme surface points with the same relative rank. As a result, fibre seeds originating at the rear of the head are connected to posterior points on the apodeme, and seeds located closer to the mandible are connected to anterior points (see SI). To prevent fibres from crossing the apodeme, the apodeme surface point closest to the fibre seed was selected from all points with the same anterior-posterior rank. The orientation of the lines connecting fibre seeds to apodeme surface was then used to calculate fibre pennation angles with respect to the apodeme main axis. The length of the connection lines *L_t_* was used to extract muscle fibre length. For directly-attached fibres, *L_d_* = *L_t_*. For filament-attached fibres, the fibre length was calculated as *L_f_* = *L_t_* - *L_fil_*, where *L_fil_* is the filament length, obtained in a last step.

Third, we extract filament length *L_fil_* for each fibre. To this end, filaments are grown from their insertion points on the apodeme along the orientation of the respective fibre. After crossing at least 5 pixels of muscle tissue, approximately equal to the fibre diameter, the growth was terminated, and the filament length extracted. Muscle fibres for which *L_fil_* < 5 pixels were classified as directly-attached; fibres shorter than twice the fibre diameter were excluded from further analysis.

In order to quantify the quality of this ranking method, we manually extracted length and pennation angle of 200 fibres. The results of these direct manual measurements were compared with the results of the ranking method by matching the corresponding fibre seeds. On average, fibres from the ranking method were 1 ± 18% shorter than those measured manually, independent of opening angle [Linear Mixed Model (LMM) with random intercepts and opening angle as fixed effect: 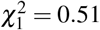, p = 0.48]; the error of the pennation angle was small, 0 ± 1°. Although this error changed significantly with opening angle – pennation angles were increasingly underestimated at larger opening angles [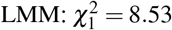, p < 0.01] – it remained small even for large opening angles, < 3°. The quality of our simple sorting algorithm thus compares favourably to that of state-of-the-art commercial tracing software [105, 112].

### Bite force measurements

In order to extract the force-length properties of the mandible closer muscle, we measured bite forces of *A. cephalotes* majors using a custom-built setup, described in detail in Püffel et al. [113]. In brief, a capacitive force sensor (maximum load: 1N, resolution: 0.002 N) is compressed when a force is applied to a bite plate connected to a pivoting lever (see Fig. 2A). The position of a second slidable lever can be controlled by a stepper motor, so adjusting the distance between two bite plates. To extract mandible and head position during biting, a camera recorded the experiment simultaneously from top-down at 30 fps. A 45 °-angled mirror provides a side view for a 3D reconstruction of landmarks (see below). Force sensor, motor and camera were controlled by a Raspberry Pi (v 3B+, Raspberry Pi Foundation, Cambridge, UK), and operated via a graphical user interface. Details on the sensor calibration are provided in the SI.

**Figure 2.**
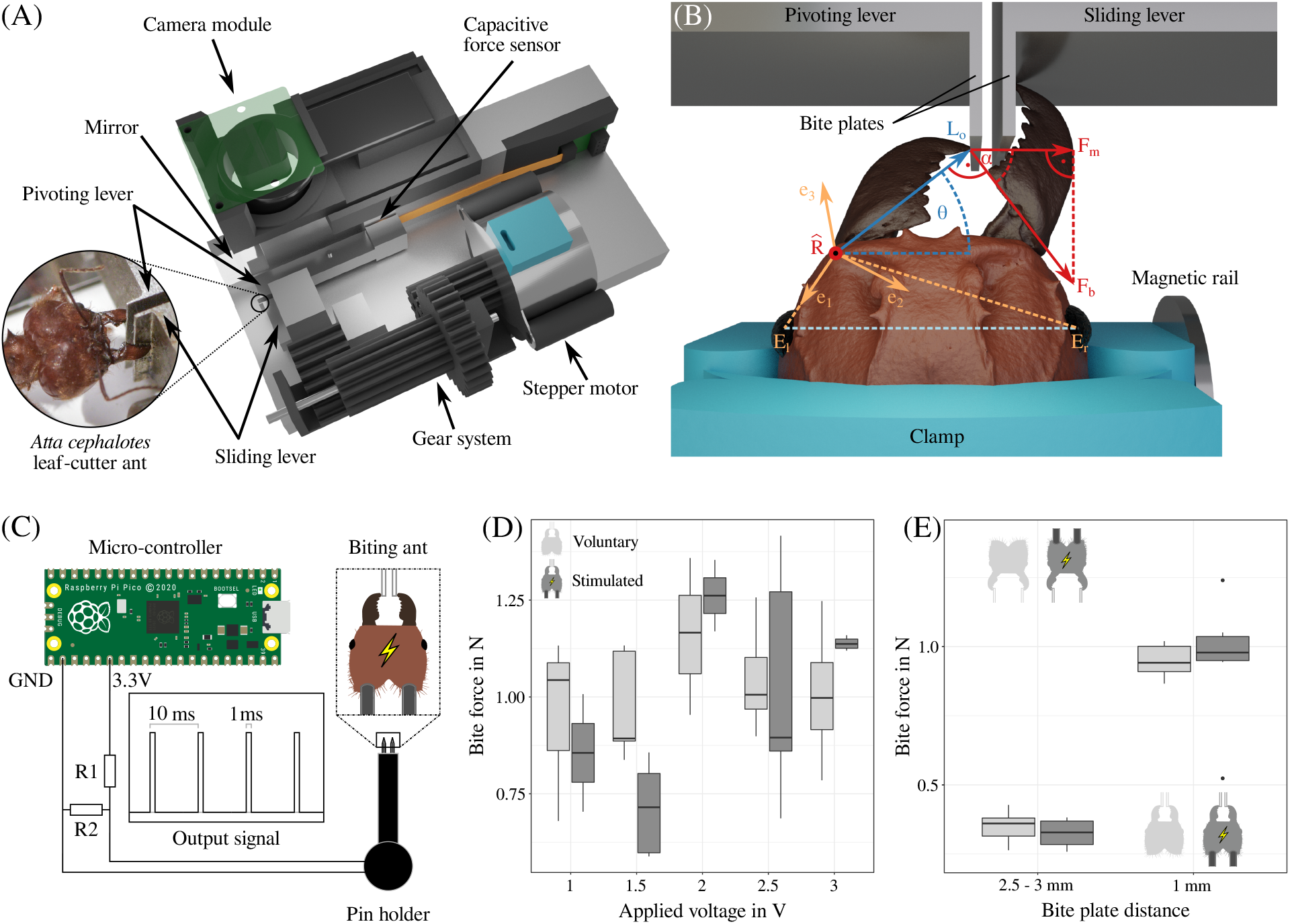
(**A**) A custom-built force rig was used to measure bite forces of *Atta cephalotes* majors (photo by Victor Kang). In short, a forcesensitive capacitor was attached to a rigid metal frame. A pivoting lever presses onto it as a force is applied at the opposite end of the lever. A second sliding lever, controlled by a stepper motor, provides control over the distance between the bite plates. (**B**) To measure bite forces at small opening angles *θ*, ants were positioned off-centre using a 3D printed clamp attached to a magnetic rail. Bite force |**F**_*b*_| was extracted from the measured force |**F_*m*_** | using the angle α, the outlever **L_*o*_** and the joint axis of rotation 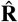, as defined in Eq. 8. 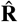 was projected onto a head-fixed coordinate system (**e**_1_, **e**_2_, **e**_3_), defined by the vectors connecting the mandible joint with left (**E**_*l*_) and right eye centres (**E**_*r*_). (**C**) A custom-built device was used to stimulate the mandible closer muscle, so eliciting maximum bite forces. A Raspberry Pi Pico was used to generate a square function as output signal with 100 Hz frequency and a duty factor of 0.1. The voltage was regulated using a voltage divider with varying resistors (0.08 ≤ R1 ≤ 2.24 *k***Ω**, R2 = 1 *k***Ω**). A 3D printed device was connected to the circuit. This device holds two insect pins, 1.5 mm apart and each protruding 2 mm. The pins were inserted into the head of live ants, positioned in front of the bite plates, and the device was activated. (**D**) Stimulated bite forces (dark grey) were compared against the maximum voluntary forces (light grey), recorded prior to stimulation for voltages between 1 and 3 V. For voltages ≥2 V, stimulated forces were approximately constant. For 1V (shaded area), bites did not always occur after activating the device; the ‘stimulated’ forces hence likely include voluntary bites. (**E**) Maximum voluntary and stimulated bite forces were of similar magnitude for both small and large opening angles (measured with a stimulation voltage of 2.75 V). Hence, ants bite with close to maximum available force independent of opening angle, and undeterred by the rigid, metallic material of the levers and the clamping.

#### Experimental protocol

Ant majors were collected from the foraging area or the fungus garden as available, and held in front of the bite plates of the force sensor using insect tweezers. Force recordings were terminated as soon as at least five distinct bites occurred. Subsequently, the ant was marked (Edding 4000 paint marker, Edding AG, Ahrensburg, Germany), and placed into a separate container with fresh bramble leaves and around 20 minors and medias to provide a resting period. After around two minutes, the same ant was measured again, but with a different plate distance. This process was repeated once more, so that bite forces of a single individual were measured three times at three different plate distances. Plate distances were chosen at random from three equally-sized bins between 0.7 and 3.6 mm, approaching the maximum mandible gape range of the ants. In order to test if muscle fatigue may confound the results [114], an additional five ants were measured using the same protocol, but at a constant plate distance of 1 mm. A LMM with random intercepts revealed no significant effect of trial number on bite forces (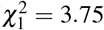, p = 0.053; we note that this result is almost significant, but in the opposite direction than expected: bite forces were around 15 % larger at the last trial compared to the initial measurement).

To comprehensively characterise the relationship between bite force and mandibular opening angle, it is necessary to measure bite forces across the largest possible range of opening angles. The upper end of this range can be approached by simply increasing the distance between the bite plates; the smallest angle, however, is structurally limited by bite plate thickness (approximately 150 *μ*m each for our set-up). To overcome this limit, we fixed the ant heads with a custom 3D printed clamp, magnetically connected to a metal frame. The clamp allowed us to position the ant heads off-centre relative to the bite plates, so enabling bites at lower opening angles (see Fig. 2B). Clamped ants did bite much less frequently, and sometimes only after stimulation with polite air blows [also see 60]. To test if clamping affected bite forces, five ants were clamped and positioned centrally in front of the sensor. The plate distance was selected at random to fall between 1.7 and 2.5 mm, and bite forces were measured. There was no significant difference in force between unclamped and clamped bites of the same gap class (Two Sample t-test: t_28_ = 1.19, p = 0.24).

#### Video analysis

We extracted the maximum force from each bite force trace (see SI for raw data examples). To avoid confounding effects due to variation in mandible outlever, only bites transmitted with the most distal tooth were considered. The following 3D coordinates were extracted from the video frame corresponding to the peak bite force: the bite contact point (most distal tooth tip), joint centre, and the geometric centres of both eyes. The depth component of the 3D coordinates was measured from the mirrored side view [113]. The vectors connecting mandible joint with both eyes **E**_*l*_ and **E**_*r*_ were used to define a local head-fixed coordinate system: **e**_1_ = **E**_*l*_/|**E**_*l*_|, **e**_3_ = (**E**_*l*_ × **E**_*r*_)/|**E**_*l*_ × **E**_*r*_| and **e**_2_ = **e**_3_ × **e**_1_. The rotational axis, 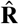, extracted from the tomographic scans, was projected onto this coordinate system using the same relative orientation to **e**_1_, **e**_2_ and **e**_3_ (see Fig. 2B). The mandibular opening angle was calculated with respect to the lateral axis, here defined by the vector connecting both eye centres.

The capacitive force sensor has only one sensitive axis, i. e. it is a 1D sensor. |**F**_*b*_| was hence extracted as defined by Eq. 8. Due to the design of the setup, an additional correction factor Γ was introduced to account for differences in moment arms due to variation in contact point on the bite plate. This correction factor reads Γ = *L_p,cal_*/*L_p,b_*; *L_p,cal_* is the moment arm used for calibration, and *L_p,b_* is the moment arm defined by the mandible contact point on the bite plate.

#### Electrical stimulation

To correctly interpret our bite force measurements, we tested (i) if voluntary bites involve maximum muscle activation, and (ii), if activation is independent of mandibular opening angle. We conducted these tests by eliciting maximum muscle activation through electrical stimulation of the mandible closer muscle.

To this end, a microcontroller (Raspberry Pi Pico, Raspberry Pi Foundation, Cambridge, UK; maximum output current 300 mA) was programmed to output a square function of stimulation impulses at 100 Hz frequency with a duty cycle of 10 % (parameters used to elicit muscle contraction in moths [115] and beetles [116, 117]). The output voltage was regulated using voltage dividers with resistors between 70 and 2230 Ω. A custom-designed device, consisting of two insect pins (size ‘0’, power and ground) and a 3D printed pin holder, was connected to the circuit (see Fig. 2C).

Unclamped bite force experiments were performed following the above protocol with a fixed bite plate distance of 1 mm. This time, however, the measurement was not stopped after a sufficient number of bites. Instead, the pins were inserted from posterior into the head capsule of the biting ant such that each pin was approximately in the centre of the closer muscle. Subsequently, the stimulation was turned on for 1 s, and the resulting bite force was recorded. Due to the approximate positioning and size of the pins, we cannot exclude that the opener muscle was also stimulated. However, due to its small relative size in *Atta* ants [≈5 % of the closer muscle in volume, see 81], any fast twitch excitation co-contraction would only have minor effects on the net bite force.

Two maxima were extracted from the recorded force trace: the maximum force of distal bites during natural and stimulated bites. Voltages between 1 and 3 V were used to find the minimum voltage at which the muscles where stimulated maximally, similar to the range used for beetle leg muscles [116]. At 1V, bites were not always elicited; the extracted second maximum may thus often represent voluntary bites. At 1.5 V the muscle may have been sub-maximally stimulated, causing lower bite forces than from the voluntary bites (see Fig. 2D). A marked increase of stimulated bite force was visible at 2 - 2.5 V, and for voltages ≥ *2V*, the average ratio between maximum voluntary and maximum elicited bite force remained constant [ANOVA: *F*_1,9_ = 1.17, *p* = 0.31]. On the basis of these results, we selected 2.75 V as excitation voltage for further stimulated bite force measurements at small (1 mm) and large (2.5 - 3 mm) plate distances. Occasionally, stimulated bites did not occur at the distal end of the mandible blade. In these cases, the calculated maximum bite force was corrected as |**F**_*b,s*_| = |**F**_*b*_||**L**_*o,c,eff*_|/|**L**_*o,d,eff*_| to account for differences in effective outlever, where **L**_*o,c,eff*_ and **L**_*o,d,eff*_ are the effective outlevers at the point of bite contact and the most distal tooth tip, respectively. We excluded stimulated measurements for which the difference in opening angle between unstimulated and stimulated bite exceeded 20 ^°^ to reduce confounding effects, and one additional measurement that exceeded the force range of sensor calibration.

#### Data analysis

The variation of morphological force determinants with mandibular opening angle can be directly predicted from the derived geometric relations (see Eqs. 2–7). As reference parameters for these predictions, we selected the corresponding arithmetic average across all scans. To test for significant effects of opening angle on a variety of morphological parameters of the bite apparatus, we deployed Linear Mixed Models: we tested if adding opening angle as fixed parameter improves the model significantly compared to a random intercept model [see 118]. To extract the force-length properties of the muscle (see Eq. 5), maximum muscle stress, optimum fibre length and shape parameter *β* were estimated using a non-linear least squares fitting function in python, invoking Eq. 1. To enforce physical and biological plausibility, boundaries were set such that the maximum muscle stress and *β* are positive; the optimum fibre length was forced to remain within the range of measured fibre lengths. The physiological properties of directly- and filament-attached fibres likely differ [see above, and 100]; however, due to the small fraction of directly-attached fibres, 15 ± 8 %, we lacked sensitivity to determine their physiological properties with sufficient confidence. The physiological muscle parameters of these fibres were thus assumed to be equal to those of the filament-attached fibres; *L_opt_* was assumed to be at the equivalent stretch with respect to the average fibre lengths of both fibre populations. We merged data from all bite force experiments, as neither clamping, nor measurement order affected bite force. We excluded the stimulated bites to avoid pseudoreplication and four further data points: three measurements from one individual that produced outliers at low opening angles (55° < θ < 65°; |**F**_*b*_| < *μ* – 3*σ*), and one measurement where the second mandible interfered during the experiment. Overall, we analysed 138 force measurements from 81 individuals (54.8±3.2mg).

## Results and discussion

Bite force influences animal behaviour and relates directly to muscle physiology and head morphology. It has significant consequences for animal fitness, and is thus under selection. As a consequence, bite forces may be considered a nonpareil performance metric for comparative studies at several levels of organismal biology [1]. We derived a comprehensive first principles model to describe the variation of bite force with mandibular opening angle across arthropods. In order to validate the geometric derivations of our model, we measured the relevant parameters from tomography scans of *A. cephalotes* majors across a large mandibular opening range. We then used our model to extract the physiological properties of the mandible closer muscle from *in-vivo* bite force measurements via Eq. 1. In the next paragraphs, we (i) demonstrate the accuracy and predictive power of the model; (ii) discuss the magnitude and variation of leaf-cutter ant bite forces with opening angle in an ecological and comparative context; and (iii) present a ‘minimal’ model to predict bite forces from simple morphological measurements.

### Variation of morphological bite force determinants is in excellent agreement with model predictions

Our model derivation involved four key assumptions: First, we assumed that filament strain is negligible. This conjecture is supported by the observation that filament length is independent of opening angle (LMM: 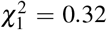, p = 0.58, see SI). Second, we assumed that the head capsule is effectively rigid. This assumption is supported by the fact that head dimensions vary little with opening angle (see SI). Third, we conjectured the apodeme ligament rotates, and so enables the lateral displacement imposed by mandible rotation. Indeed, we report evidence for such changes in the lateral expansion of the ligament in the SI. Fourth, we modelled the mandible joint as a one degree-of-freedom hinge. The mandible joint in some ants, including leaf-cutter ants, has an unusual morphology [119], but mandible motion nevertheless approximates planar rotation across large change in opening angle relevant to this study [Kang, Püffel and Labonte, in preparation].

Having provided empirical support for the simplifying assumptions involved in the model derivation, we turn our attention to the predictions it enables. Mechanical advantage, apodeme displacement, fibre pennation angle and fibre length are important morphological determinants of bite forces, which all change with mandibular opening angle (see Eqs. 2–7). Direct morphological measurements of these parameters across a large range of opening angles are in excellent agreement with theoretical predictions based on arithmetic means, without exception (see Fig. 3).

**Figure 3.**
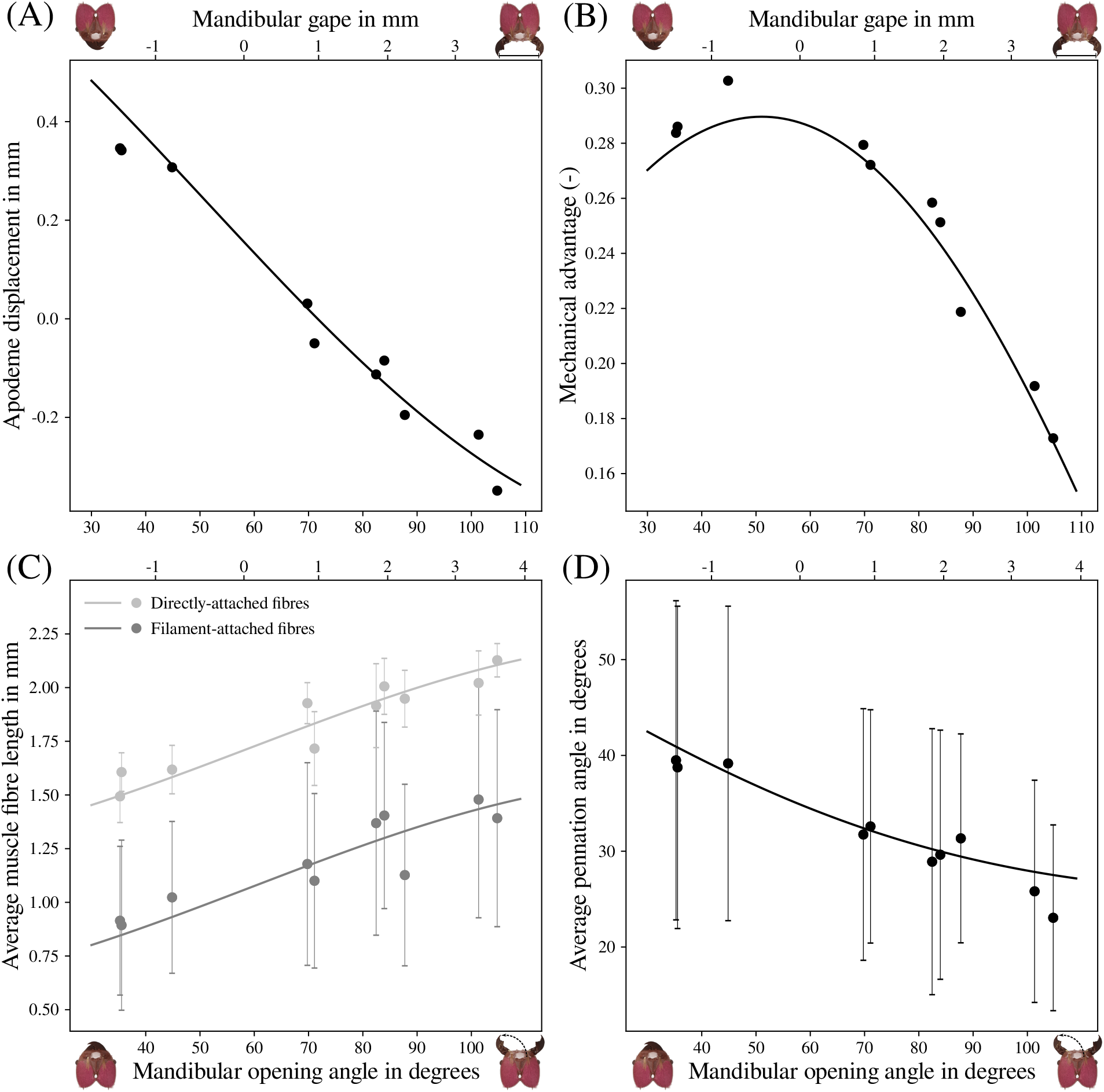
The variation of the relevant morphological parameters with mandibular opening angle, extracted from tomographic scans of *Atta cephalotes* majors, is in excellent agreement with the theoretical predictions. Points and error bars represent the extracted mean±standard deviation. The solid lines show the model predictions based on the sample means (see text); they are not fitted lines. The mandible gape is shown on the upper abscissa. (**A**) Mandible closure is associated with a displacement of the apodeme. Across the opening range, this displacement amounts to around 0.7 mm, or about one-fifth of the average head length. Positive values represent displacements away from the average apodeme position in the posterior direction, and negative values indicate anterior displacements relative to the average position. (**B**) The mechanical advantage is maximal at 51°, close to zero mandible gape, and decreases by a factor of almost two at the largest opening angle. (**C**) The displacement of the apodeme is associated with a change in fibre length, which follows different relationships for directly- and filament-attached fibres (see Eqs. 6 and 7 in the text). The resulting *absolute* length changes are similar between both fibre populations (≈ 0.6mm). However, as directly-attached fibres are much longer on average, their *relative* length change is much smaller. (**D**) As fibres shorten, they rotate and thus change their orientation. Across the opening range, the associated decrease in average pennation angle is around 13°.

We measured opening angles between 35 and 105° for fully closed and maximally opened mandibles, respectively. At the maximum opening angle, the gape is 3.6 mm, or approximately 75 % of the head width. At the minimum gape, in turn, the most distal mandible teeth overlap by about 1.3 mm. To move the mandible from maximally opened to fully closed, the apodeme displaces by around 0.7 mm corresponding to about one-fifth of the average head length (see Fig. 3A). This displacement may appear small, but it constitutes more than 50% of the average fibre length, and thus requires large fibre strains (see below). At small opening angles, the apodeme angle, *γ* is close to 90°, and varies approximately linearly with apodeme translation (this follows from the small angle approximation, cos(*dθ* + 90°) = −sin(*dθ*) ≈ −dθ, see Eq. 3). At large opening angles, this linearity no longer holds, and the same change in opening angle is associated with a smaller longitudinal displacement of the apodeme attachment point, and a larger lateral displacement (see Fig. 1E).

The mechanical advantage takes a maximum value of 0.29 at an opening angle of 51°, where *γ* ≈ 90°. As a consequence, at most 29 % of the apodeme force is transmitted at the distal end of the mandible blade (see Fig. 3B). The maximum *MA* is at the lower end of values reported for insects across numerous taxa [0.3 < *MA* < 0.8, see 84]. The *MA* decreases with increasing opening angle in direct proportion to the effective inlever, and is reduced by a factor of 1.7 at the largest opening angle, at which *γ* has decreased to about 40°. The increasing steepness of this decline directly reflects the change in slope of sin(*γ*) (see Eq. 2). If the variation of *MA* was neglected, *MA*(*θ*) = *MA_max_*, the capacity for bite forces at large opening angles would thus be overestimated by about 70 %. This variation is similar to the range of possible effective outlevers from the most distal to the most proximal sections of the mandibular blade for our study organism and across various insect taxa [84]. The fact that animals can control *MA* by changing the outlever has previously been discussed in an evolutionary context [84]; that a similar effect, which alters bite force just as much across the opening range, arises from variations in opening angle, however, has received very little attention in both the vertebrate and invertebrate literature [but see 80, 82].

The displacement of the apodeme is associated with a change in fibre length. Both directly-and filament-attached fibres shorten by around 0.60mm across the opening range, starting from a similar maximum *total* length of 2.11 and 2.07 mm, respectively (see Fig. 3C). The absolute changes in fibre length are similar, but with an average filament length of 0.60 mm – equivalent to around 75 % of the smallest fibre length – the relative changes in fibre length differ substantially. The ratio between largest and smallest fibre length is 1.41 for directly-attached fibres, but 1.73 for filament-attached fibres, considerably larger than ratios reported for other insects (1.35 and 1.55 for the mandible closer muscle in stag beetles and cockroaches, respectively, [10, 82], and 1.50 for the extensor tibiae muscle in stick insects [120]). Such large changes in relative muscle length are associated with a substantial decrease in muscle stress [93, 94, and see below], which seemingly favours a direct muscle attachment over filament-attachment. The abundance of directly-attached fibres, however, is likely limited by space constraints in the head capsule; filament-attachment increases the effective internal muscle attachment area to the apodeme, so that its net effect may still be an increase in muscle force [81, 103].

In addition to changes in fibre length, the apodeme displacement is associated with changes in average pennation angle (see Fig. 3D). *φ* is around 41^°^ at low opening angles and decreases to 27^°^ at large angles. The notable standard deviation of the measured angles (and fibre lengths, see Fig. 3C) does not reflect measurement error (see method validation), but the multi-pennate architecture of the closer muscle in *Atta* ants [see 81, 100]. The decrease of φ is steepest at small opening angles, and flattens towards larger angles (see Fig. 3D). This pattern is driven by two effects: First, at large opening angles, the apodeme attachment point displaces mostly laterally, due to the small apodeme angle. As a consequence, the longitudinal displacement, which controls fibre rotation, is small. Second, the change in pennation angle per unit apodeme displacement is determined by the magnitude of the pennation angle itself, and is largest for large pennation angles.

The decrease in *φ* increases the fraction of the muscle force aligned with the apodeme by about 20 %, cos(27°)/ cos(41°) ≈ 1.18, more than twice as much as reported for cockroaches [82]. This increase is nevertheless small, and insufficient to balance the reduction in force due to the decrease in *MA* (Fig. 3B). There exists a further notable difference between the two parameters: the pennation angle will always be maximal for small opening angles and then decrease monotonously to a minimum at large opening angles (see Fig. 1). In sharp contrast, the effective inlever is largest at *γ* = 90°, which could be achieved at any opening angle by relatively small changes in mandible shape, so resulting in non-monotonous variation across the opening angle range. These observations suggest two different functional roles for the pennation and apodeme angle: The variation in pennation angle is likely irrelevant for the change in force magnitude across opening angles. However, large pennation angles increase the maximum possible physiological cross-sectional area in the limited volume of the rigid head capsule, and thus control the total magnitude of the bite force [53, 81, 103]. Pennation also results in ‘displacement’ gearing [see 121, 122], which reduces the fibre strain required to cover the same range of opening angles. This reduction is important, because muscle stress may decay steeply with fibre length; it is however difficult to draw direct conclusions from it, because the interactions between pennation angle, fibre strain, muscle stress, and the maximum possible muscle size in a closed volume are subtle and require a more detailed parametric analysis than is within the scope of this study. The apodeme angle, in turn, does not meaningfully alter the maximum magnitude of the bite force. However, because it can increase or decrease with opening angle, it can be used to counter or magnify any changes in bite force arising from force-length effects – it is a more flexible gearing parameter across the opening range than the pennation angle. The possible ‘design space’ span by these two parameters is thus large, and presents an interesting area for comparative work.

We have demonstrated that the variation of the key morphological design parameters of the insect bite apparatus with opening angle can be accurately predicted from first principles. Next, we show that the knowledge of these relationships can be used to extract the physiological properties of insect muscle, which remain understudied [but see 100, 120].

### Leaf-cutter ants are extremely specialised for large bite forces, and bite strongest at opening angles relevant for cutting

We used the deductive power of our geometrically validated biomechanical model to extract the force-length properties of the mandible closer muscle. To this end, we performed *in-vivo* bite force measurements across a large range of opening angles, 50-105^°^, and extracted the magnitude of the maximum bite force at each opening angle via Eq. 8. To test if the ants bit with maximum activation, we compared voluntary and electrically stimulated bite forces. There was no significant difference between the maximum voluntary and stimulated bite forces for neither small nor large bite plate distances [Welch Two Sample t-test, 1mm distance: t_5.68_ = −0.04, p = 0.97; 2.5-3 mm distance: t_9.69_ = −1.76, p = 0.11, see Fig.2E]. Consequently, ants appear to maximally activate their muscle during our bite force measurements, independent of the mandibular opening angle, and undeterred by the rigid material of the bite plates [but see 123].

In order to characterise the physiology of the mandible closer muscle, we extracted maximum muscle stress, optimum fibre length, and the force-length shape parameter *β*, via a nonlinear least squares fit of Eq. 1. The resulting fit yielded optimum fibre lengths of *L_opt_* = 1.42mm [95 % CI (1.17; 1.66)] for directly-attached fibres, and *L_opt_* = 0.92mm [95 % CI (0.79; 1.07)] for filament-attached fibres. We used *L_opt_* to extract the physiological cross-sectional area of the muscle at a defined point on the force-length curve (see above), *A_phys_* = *V_*m*_*/*L_opt_* = 4.3 ± 0.4mm^2^. The maximum muscle stress was fitted as 1.16MPa [95 % CI (1.06; 1.26)], at the upper end of values reported for arthropod muscle, which are typically below 1 MPa [10, 53, 60, 62, 120, 124, but see 16, 23]. We caution against strong conclusions on the basis of this comparison, because most of the published muscle stress estimates do not represent the ‘true’ maximum muscle stress, as they do not stem from direct or indirect measurements of force-length properties. Based on the difference in average sarcomere length, it is likely that the true stress of some crab pincher muscles, for example, exceeds our estimate for leaf-cutter ant closer muscle [see 23, 100], which may suffer further from shrinkage effects arising from CT sample preparation [125].

The shape parameter was fitted as *β* = 5.34 [95 % CI (0.09; 10.59)]. Data on the force-length properties of arthropod muscle are surprisingly scarce. In order to assess the plausibility of our result, we extracted force-length data from published work, normalised all forces with maximum force, all lengths with optimum length and fitted Eq. 5 with a non-linear leastsquares algorithm. The shape parameter for *Atta* closer muscles is close to the shape parameter for fibres from the extensor carpopoditi muscle of the European crayfish [*β* = 6.27, 95% CI (5.72; 6.84), 126], the extensor tibiae muscle of Indian stick insects [*β* = 3.95, 95% CI (3.29; 4.71), 120], fibres from the meropodite muscle of walking legs of Orconectes crayfish [*β* = 6.61, 95% CI (5.87; 7.41), 127], and the abdominal ventral superficial muscle of Hermit crabs [*β* = 6.46, 95% CI (5.25; 7.87), 128]. However, it is substantially smaller than values for extensor muscles in the coxa of discoid cockroaches [*β* = 12.18,95% CI (11.10; 13.30), 129], the respiratory muscle in green crabs [*β* = 26.66,95% CI (13.07; 56.23), 130], and the (synchronous) mesothoracic dorso-longitudinal muscle of *Manduca sexta* moths [β = 58.65, 95% CI (52.89; 64.59), 131]. In light of the considerable variation in both sarcomere lengths and the ratio between thick and thin filaments in arthropod muscle [23, 132], this large range may not be surprising, and suggests exciting opportunities for comparative muscle physiology.

Our biomechanical model captures the general shape of the variation of bite force with opening angle both quantitatively and qualitatively. The agreement between prediction and observation, however, is arguably less convincing than for the geometric relations alone in at least two aspects (see Fig. 4A). First, at the largest opening angles, the fit systematically underestimates the measured bite forces, which remain approximately constant for *θ* > 90°. This disagreement partially reflects passive joint or muscle forces, which may contribute about 50 mN [see SI, and 60, 101]. Second, close to the fitted maximum, the spread of measured bite forces is large. At small opening angles, the misalignment angle between measurable and bite force vector is most extreme (see Eq. 8). The increased variation may thus represent a combination of biological variation and small errors in landmark placement, amplified by larger correction factors. For measurements at small opening angles (< 65°), the average correction was 28 ± 25 %, compared to only 5 ± 4 % for measurements at larger angles.

**Figure 4.**
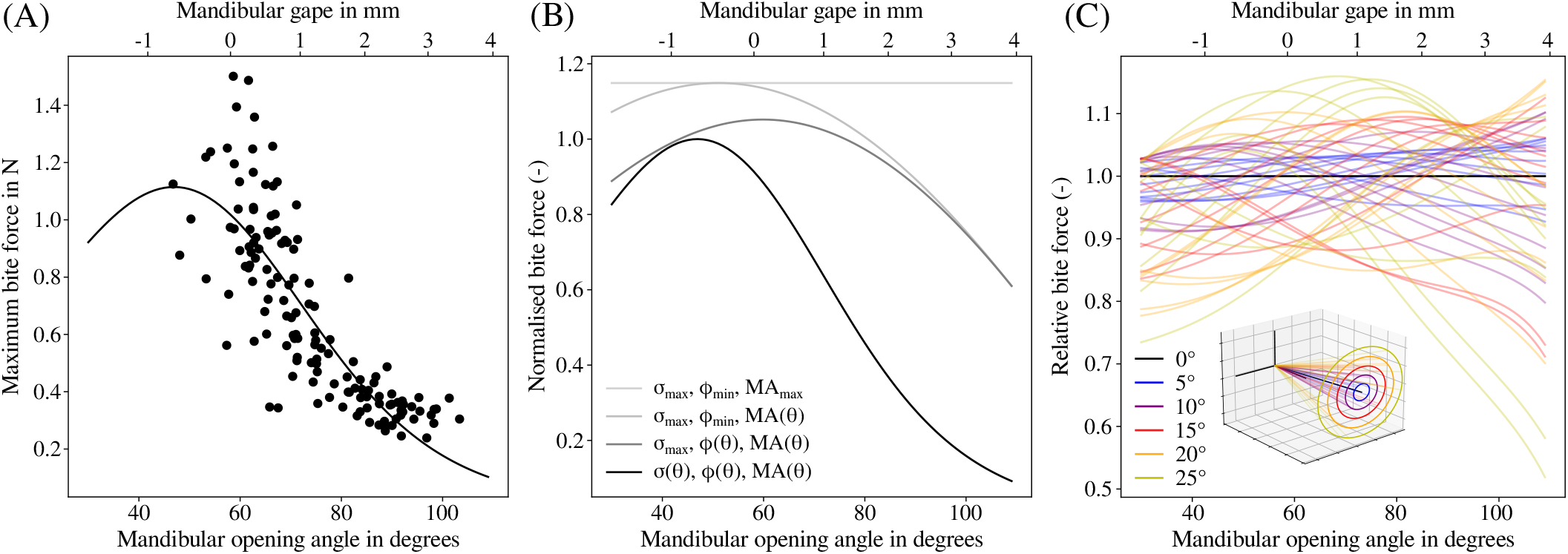
(**A**) Bite forces of *Atta cephalotes* majors were measured across mandibular opening angles between 50 and 105°. Bite forces peak at small opening angles, and decreased to around one-fifth of that maximum at 100°. The solid line is a least-squares fit of Eq. 1 which accounts for the variation of both morphological and physiological determinants of bite force. For the largest opening angles, the model underestimates the measured forces, which may be partially attributed to passive joint and muscle forces (see SI). The peak muscle force associated with peak bite forces is 9,000 times larger than the mean body weight of the ants – a factor of seven larger than expected for animals of this size (see text). (**B**) The variation in bite force is driven by changes in muscle stress and mechanical advantage. The pennation angle, on the other hand, mainly modulates the maximum bite force magnitude. The black line represent the fitted relationship between bite force magnitude and opening angle, |**F**_*b*_|(*θ*), normalised with its maximum. The grey lines show the theoretical relationships between bite force and opening angle, if muscle stress, the cosine of the pennation angle, and the mechanical advantage were progressively fixed at constant values equal to their respective maxima. If these parameters were all maximal at the same opening angle, the maximum bite force would increase by a mere 15 %. Because muscle volume occupancy is also close to its maximal theoretical value [81], leaf-cutter ant heads are close to a morphological optimum for maximum bite forces. (**C**) The variation of bite force with opening angle strongly depends on the mandible joint axis of rotation. A seemingly small variation in this axis of only 15°, changes the relative bite force prediction by up to 30 %. The coloured lines represent the theoretical relationships between bite force and opening angle for 50 randomly generated axes of varying angles to the actual axis (black), normalised with the fitted relationship, |**F**_*b*_|(*θ*). Clearly, the axis of rotation is an absolutely crucial parameter if the variation of bite force with opening angle is to be predicted accurately.

We measured maximum bite forces of 1.4 N, equivalent to about 2600 times of the mean body weight of the ant workers. Maximum bite forces were measured at small opening angles (< 65°, see Fig. 4A), and bite force decreased steeply with opening angle to a minimum of 0.3 N at 100° - a total variation of a factor of five. Even considering the comparatively small size of the ants (50-60 mg), these weight-specific forces are rather remarkable indeed: Alexander conducted a literature review of the maximum forces exerted during various activities, including biting, by animals spanning ten orders of magnitude in mass [79]. He reported an upper bound for weight-specific forces of *F_max_* = 20m^-1/3^, where *m* is the body mass in kg. The expected value for a 55 mg leaf-cutter ant is *F_max_* = 20 (55 10^-6^)^-1/3^ ≈ 500. Leaf-cutter ants thus bite with a force around five times larger than expected for an animal of their size; these values even surpass those from other unusually strong animals such as crabs [23]. The specialisation of leaf-cutter ants to produce large bite forces is even more startling in the context of muscle force: Leaf-cutter ants exert maximum muscle forces of *σ_m_A_phys_* ≈ 5N – around 9000 times their body weight. A typical upper bound for maximum weight-specific muscle force is 50m^-1/3^ [79], or ≈ 1300 for leaf-cutter ant majors – almost an order of magnitude smaller than the observed value. Clearly, leaf-cutter ants are extremely specialised to produce large bite forces.

Having fully characterised peak bite forces, and the key physiological and morphological parameters of the mandible closer muscle in *Atta* majors, we can assess the relative importance of physiology or morphology for the bite force-gape relationship (see Eq. 1). The majority of this variation is driven by changes in muscle stress, which varies by a factor of 5.7 (see Fig. 4B); the mechanical advantage decreases by a factor of 1.7. These effects are slightly attenuated by the decrease in pennation angle, which effectively increases the bite force for large opening angle by 18 %. Remarkably, if all parameters were optimum at the same opening angle, the maximum bite force would increase by only ≈ 15 %. Compared to the significant variation in muscle stress and mechanical advantage, this ‘lost’ potential appears rather small, and it is mainly driven by the pennation angle, which is anatomically constrained (see above). The bite apparatus in leaf-cutter ants thus seems to have an architecture close to a theoretical optimum; the muscle volume density approaches its theoretical maximum [81], and pennation angle, apodeme angle and optimum fibre length are all ‘synchronised’ such that bite force peaks in a narrow range of opening angles (centred around 47°). The opening angle range where bite forces are maximal corresponds to small mandible gapes (< 0.5 mm). This range is most critical in the context of the typical behaviour displayed during leaf-cutting, because fresh cuts are often initiated with ‘scissor-like’ cuts through the leaf lamina, and the average lamina thickness of tropical leaves is around 0.25 mm [133]. When propagating cuts through the lamina, ants often use a large range of opening angles [134]; however when encountering thick veins that require high bite forces, mandibles open only very little [see SI in 135]. Leaf-cutter ants may thus adjust their cutting behaviour in accordance with the steep decrease of bite force for opening angles > 75°. These results represent a key step towards understanding the intricate relationship between cutting behaviour, maximum bite force and mandibular opening angle in *Atta* foraging [also see 113].

### Minimal models for the magnitude of bite force and its variation with gape

We have demonstrated that both the magnitude of bite force and its variation with gape can be accurately predicted from first principles. Many individual components of our biomechanical model have been discussed by colleagues previously [10, 53, 56, 60, 80, 81, 136–138, e.g.]. However, a comprehensive analysis which combines all elements into one single model has to our knowledge been absent from the literature. The advantage of a complete first principle model is that it is as accurate as the estimates of the relevant parameters which define it. The problem is that many of these parameters are time-consuming and costly to measure. Accordingly, there has been considerable interest in ‘minimal’ bite force models, where some parameters are either replaced with first order approximations, scaling relationships, or proxies identified via statistical analysis [1, 3, 26, 28, 56, 136, 138–141]. As an illustrative example, head width has repeatedly been shown to correlate tightly with bite force in vertebrates, which, combined with conserved maximum muscle stress, seemingly makes bite force predictions rather straight forward [e. g. 6, 28, 138]. Due the extraordinary attractiveness of a minimal bite force model for ecologists, palaeontologists, evolutionary biologists and biomechanists alike, we next discuss the implications of our analysis for such models in insects; first with respect to the magnitude of the absolute bite force, and then for its variation with gape. In probing the accuracy with which the magnitude of the maximum bite force can be predicted without full knowledge of all parameters, we are assessing an upper bound, corresponding to the assumption that all relevant parameters take their maximum value at the same opening angle (see Fig. 4B). By separating the question of bite force variation with gape from its absolute value, we are then considering only *relative* changes of bite force with gape.

The maximum distal mechanical advantage of the mandible bite apparatus in insects varies between 0.3 - 0.8 across a broad range of insect orders [84]. Using a value in the middle range of *MA* ≈ 0.55 is thus associated with an uncertainty of less than a factor of two. A more accurate proxy may be obtained by measuring the ratio between mandible width and length [see e. g. 53]. The mandible length is likely a relatively accurate proxy for the outlever, because the uncertainty associated with the location of the joint centre will be small compared to its length. The mandible width, however, may be a poor proxy for the inlever, which is typically shorter than the outlever, but subject to the same absolute uncertainty.

Average pennation angles of mandible closer muscles in insects are smaller than 45° [data exists for ants, beetles, and cockroaches 81, 82, 103, 142]. The direct influence of pennation on bite force magnitude is thus negligible, cos φ ≈ 1.

An accurate measure of *A_phys_* is perhaps most challenging to obtain: it requires knowledge of muscle volume, and crucially, the optimal fibre length. *L_opt_* is typically unknown, and can only be estimated from TEM images of muscle tissue (to measure thick and thin filament length and lattice spacing), or via knowledge of all other parameters and bite force measurements (this study). Using the physical cross-sectional area of muscle at an arbitrary but naturally occurring muscle length is associated with an error directly related to the strain range of the muscle: because muscle is incompressible, any change in its length is associated with a change in its cross-sectional area such that volume is conserved, *A* = *V/L*. If *L_opt_* was overestimated by 70 %, corresponding to the maximum fibre length ratio extracted in this study, *A_phys_* would be underestimated by 1 – 1/1.7 ≈ 40 %. However, even this proxy requires knowledge of muscle volume and fibre length, and thus CT data and advanced fibre tracing algorithms [81]. Due to these difficulties, a large number of proxies have been used in the literature, including head size (see below), the surface area of the apodeme [14], or the muscle attachment area to the head capsule [10]. Head size is arguably the most attractive proxy for *A phys*, for it represent the best compromise between accuracy and ease of measure; head volume can be readily estimated from head length, width and height, *V_h_* ≈ *H_w_H_h_H_l_*, with simple light microscopy. In insects, both mandible closer muscles typically occupy between 10 and 50 % of the total head volume *V_h_* [81, 82, 142,143, but see 104]. The volume of a single closer muscle can thus be estimated as *V_m_* ≈ 0.3/2*H_w_H_h_H_l_*. *L_opt_*, in turn, is some fraction of a linear head dimension. The most suitable dimension may be head height, as the region where it takes its maximum value is mainly occupied by closer muscle [see 81, 82,142, 143]. It follows that *L_opt_* ≈ 0.5*H_h_* and *A_phys_* = *V_m_*/*L_opt_* ≈ 0.3*H_w_H_l_*. This estimate has an error similar to the suggested approximation for the mechanical advantage, and is as good as the implicit yet untested assumption of isometry across animals of wildly different sizes.

The maximum muscle stress is the parameter most prone to variation; estimates for arthropods vary by almost two orders of magnitude [0.04 - 2.2MPa, see 16, 23, 60, 120, 124]. This large range limits the accuracy of a mean stress to one order of magnitude at best [60], and is likely at least partially owed to the experimental methods with which stress has been estimated. Force measurements with isolated insect muscle let alone single fibres are rare [e.g. 120, 124, 129]. Most often, stress is estimated from more integrated bite/pinch force measurements instead [e. g. 23, 53, 60, and this study]. Any such estimation of muscle stress, however, is subject to the uncertainties in determining *MA* and *A_phys_* discussed above. A more accurate peak muscle stress estimate may be obtained from the average fibre sarcomere length, *S_l_* [23, but see 47]. To this end, we extracted all non-vertebrate data from the classic work of Taylor, and conducted a standardised major axis regression, *σ* ~ *S_l_*, in log-log space. For sacromere lengths between 3 - 17*μ*m, this relationship reads *σ_max_* = 50*S_l_*, where *S_l_* is in *μ*m and *σ_max_* is in kPa, which explains about 75% of the variation in stress with sarcomere length. The remaining variation is likely attributed to variations of the relative fibre length at which forces were measured and generic measurement error. The scaling coefficient of unity is robust (and has a theoretical foundation), but the proportionality constant of 50 kPa μm^-1^ is again subject to uncertainty in *A_phys_*, required to convert the measured force to a stress (for example, our measurements in *Atta* are a noteworthy outlier). Taylor estimated *A_phys_* as the surface area of the apodeme multiplied with sin(*φ*) [23, also see 15, 53]. The error in measuring apodeme surface area is likely small, but the error in assuming that it is equal to the muscle attachment area is ≥ 10% (the maximum packing density of a lattice of circles). The total uncertainty associated with this approximation for stress is thus at least 35%. Although measuring sarcomere length is technically involved, the significant reduction in uncertainty for the maximum stress associated with it may make these measurements well worthwhile.

We conclude the discussion for maximum bite force by summarising that the simplest reasonable estimate for the maximum bite force requires only knowledge of head width and length. However, this estimate may at worst only be accurate to less than an order of magnitude. The next best estimation replaces the maximum stress with a proxy based on sarcomere length, and reads:

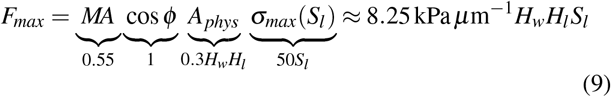

If we use an intermediate sarcomere length of 10 *μ*m [23], Eq. 9 simplifies to *F_max_* ≈ 83kPa*H_w_H_l_*. On average, this upper bound is approximately three times larger than values measured for 653 insect species [assuming 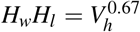, see 63]. Notably, these force measurements were conducted at opening angles which may differ from the angle of maximum force. As demonstrated above, large deviations from this angle may lead to substantially lower bite forces (factor of up to five). For 12 % of the measured species, our prediction underestimates the maximum bite force, which likely reflects species-specific adaptations. Overall, the ratio between maximum predicted and measured bite force ranges from 0.30 to 27, across almost two orders of magnitude. The effort required to measure sarcomere length is likely rewarded with a reduction in such worst-case errors by at least a factor of five.

Next, we address the variation of bite force with gape, which has received even less attention than maximum bite force, but is of similar ecological significance [e. g. 26, 80]. The cosine of the average pennation angle will typically vary at most between unity and 0.7 between maximally open and fully closed mandibles, respectively (cos0° and cos45°, see above); this variation is small enough that it can be neglected, cos[*φ*(*θ*)] ≈ constant. Instead, the variation of bite force with opening angle is likely dominated by changes in muscle stress due to changes in muscle length; such effects are particularly pronounced when fibre strain and shape parameter *β* are large, and *L_opt_* occurs at fully open or closed mandibles (this study). *β* appears to cluster around ≈ 6, but may be twice as large in some muscle (see above). The variation of the mechanical advantage can be significant, but much of this variation is governed by the apodeme angle *γ*, which can be measured relatively easily. Perhaps the most important *and* most often neglected source of variation comes from the joint rotational axis. 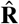 affects virtually all bite force determinants, apart from *Aphys*, and also influences the accuracy of bite force measurements themselves (see Eq. 8). In order to gain some intuition for both the error and the variation of force-gape curves which can be associated with variations in 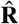, we calculated the relationship between |**F**_*b*_| and opening angle using a set of 50 alternative rotational axes, located at the same joint centre, but oriented at an angle between 5 and 25^°^ relative to the original 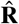 (all other parameters were kept constant). The resulting variation, normalised with the original relationship |**F**_*b*_|(*θ*), is substantial, and includes a large range of different force-gape relationships (see Fig. 4C). As illustrative examples, an axis deviating by only 10° alters the force prediction by up to 20% compared to the original relationship |**F**_*b*_|(*θ*); an axis deviating by 25° causes changes of up to 50%. Misidentifying the joint centre would lead to additional significant variation affecting both levers, the predicted apodeme displacement, and all associated parameters. For species with two well-developed condyles, the location and orientation of 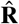 may be identified with reasonable accuracy from joint morphology alone, as previously done for beetles [10, 142], cockroaches [82] and dragonflies [61]. Other insects such as some hymenopterans, however, have more complex mandible joints [144,145], which may also have more than one degree of freedom [89, 146, 147], rendering such deductions difficult [but see 119, Kang, Püffel and Labonte, in preparation]. Given the sensitivity of bite forces to the joint axis, it is surprising how little attention has been paid to rigorously determine joint kinematics across the insect tree of life using the tools of rigid body mechanics [but see 146, and Kang, Püffel and Labonte, in preparation]. Any attempt to predict or measure the bite force-gape relationship needs to take due note of the importance of 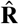 in order to avoid significant errors.

## Conclusion and outlook

We presented a validated model which predicts the bite force-gape relationship in arthropods from first principles, based on a minimum set of assumptions. We hope that our work will be useful for future studies on the bite performance in arthropods in at least three aspects. First, it enables *in-vivo* extraction of force-length properties from bite force measurements, which remain scarcely reported in the literature. The significant variation of the physiological make-up of insect muscle suggests exciting avenues for comparative muscle physiologists, and our model allows colleagues to side-step challenging measurements with tiny isolated muscle fibres. Second, our model identifies the key determinants of maximal bite forces, including a ‘minimal’ bite force model, and so facilitates comparative analyses of the functional morphology of the insect bite apparatus, including rare, small or extinct species, for which bite force measurements may be impossible. Third, it provides clear guidance on how muscle morphology and physiology translate into a variation of bite force with gape, which is of significant ecological and behavioural relevance, for example in the analysis of feeding guilds or niche formation. Indeed, the increasing number of high-resolution CT scans of insect heads offers tantalising opportunities to study the bite force-gape relationship across the insect tree of life without the need for direct bite force measurements. Notably, systems that operate close to the maximum possible bite force, such as in leaf-cutter ants, will suffer from a steeper decrease in bite force at opening angles departing from the optimum, suggesting that the functional morphology of the bite apparatus is subject to a trade-off. All three points will contribute to our general understanding of the performance, behaviour, ecology and evolution of arthropods.

## Acknowledgments

We would like to thank Victor Kang for his experimental work on the mandible kinematics of *A. cephalotes* majors, Flavio Roces for discussions and general support, and Natalie Holt and Mattia Bacca for discussions and input on the force-length relationship of arthropod muscle. This study is part of a project that has received funding from the European Research Council (ERC) under the European Union’s Horizon 2020 research and innovation programme (Grant agreement No. 851705) and a Human Frontier Science Programme Young Investigator Award (RGY0073/2020) to DL. The *μ*CT work was supported by the Advanced Imaging Materials (AIM) core facility (EPSCR Grant No. EP/M028267/1), and the European Social Fund (ESF) through the European Union Convergence programme administered by the Welsh Government (80708).

## Supplementary Materials

### Model derivations

As the mandible opens and closes, the apodeme displaces along its main axis by an amount Δ with respect to a reference position. We assume the muscle expansion perpendicular to the apodeme main axis to remain constant (i. e. the head capsule to be rigid), so that the muscle fibres rotate around their origins at the head capsule [see 102, and Fig. 1D in the main text]. As a consequence, a fibre with a total length *L*_*t*,0_ at a pennation angle *φ*_0_ shortens to a length *L_t_* at an angle *φ*, where the expansion of the fibre along the lateral axis remains constant as sin *φ*_0_***L***_*t*,0_. The fibre expansion along the apodeme main axis, however, decreases from cos *φ*_0_*L*_*t*,0_ to cos *φ*_0_*L*_*t*,0_ - Δ. As a result, the pennation angle *φ* changes as:

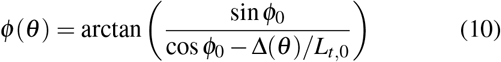

equivalent to Eq. 4 in the main text. For directly-attached fibres, the fibre length is equal to the total length, *L_d_* = *L_t_*, and follows directly from the Pythagorean theorem:

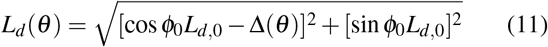

equivalent to Eq. 6 in the main text. For filament-attached fibres, the fibre length is equal to the total length minus filament length, *L_f_* = *L_t_* – *L_fil_*; the relationship between fibre length and apodeme displacement is hence modified, as:

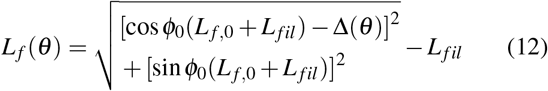

equivalent to Eq. 7 in the main text. Key here is that the fraction of the total length occupied by the filament is of a constant length (see below). As a result, the fibre strain for a given Δ is larger for filament-attached fibres compared to directly-attached fibres of the same total length.

The magnitude of bite force |**F**_*b*_| was derived as a function of mandibular opening angle (see Eq. 1), because this relationship is biologically relevant, and it allows us to compare our results to existing literature [e.g. 10, 26, 60]. |**F**_*b*_| can, however, also be expressed as a function of fibre stretch, in accordance with the causal relationship:

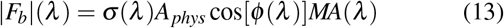

The change of muscle stress follows directly as:

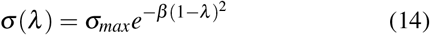

The fibre length is equal to the product between λ and the optimum fibre length *L_opt_*. Consider two fibres, one directly- and one filament-attached, of the same total length. If the apodeme displaces to the same Δ, their pennation angles change both as defined in Eq. 4 in the main text. If both fibres shorten to the same stretch ratio, however, their change in pennation angle differs. For directly-attached fibres, *φ_d_* is determined by a reference pennation angle *φ*_0_ at a fibre length *L*_*d*,0_ = *λ*_0_*L_opt_* and *λ*. In analogy to the previous derivation of *φ*(*θ*), the lateral expansion of the fibre is assumed to be constant, sin *φ_d_λL_opt_* = sin *φ_0_λ_0_L_opt_*, resulting in:

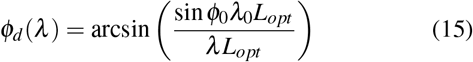

Correspondingly, the pennation angle for filament-attached fibres changes as:

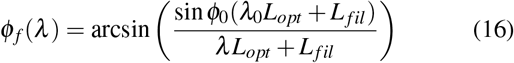

For simplicity, the relationship between apodeme displacement and effective inlever with *λ* will be derived for directly-attached fibre only. The corresponding relationship for filament-attached fibres, however, follows the same geometric relations.

The apodeme displacement Δ increases with a decrease of fibre expansion along the apodeme main axis as:

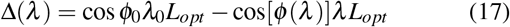

The effective inlever is determined by the length of the inlever projection onto the plane of rotation, 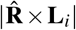, the fraction of the apodeme main axis that lies in this plane 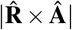, and the starting configuration between inlever and apodeme, characterised by the apodeme angle *γ*_0_ measured at a reference stretch ratio *λ*_0_ (see Fig. 1E in the main text). At *λ*_0_, the effective inlever is equal to 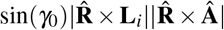, equivalent to Eq. 2 in the main text. Following the Pythagorean theorem, this length can also be expressed as 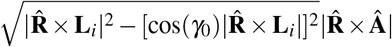, where the term 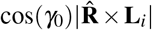 captures the expansion of the inlever along the apodeme main axis in the plane of rotation. This expansion decreases as Δ increases, attenuated by the misalignment between apodeme main axis and rotational plane. As a result, the effective inlever varies with *λ* as:

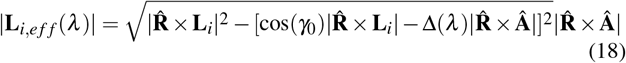

### Filament strain, head deformation and the rotation of the apodeme ligament

We made three assumptions on the mechanical properties of the bite apparatus. First, we assumed that filament strain during muscle contraction is negligible. In order to estimate filament strain, we invoke Hooke’s law, *ε* = *σ_fil_*/*E_fil_*, where *σ_fil_* is equal to the maximum muscle stress times the ratio between the crosssectional areas of fibre and filament. The fibre cross-sectional area was extracted from the scans described in this study, the filament cross-sectional area was estimated from synchrotronbased scans with a higher resolution performed on *Atta vollen-weideri* majors, described in detail in [81]. The Young’s modulus of the filaments *E_fil_* was assumed to be equal to that of the head capsule (≈ 6 GPa, F. Pueffel, unpublished data), because their density appeared to be similar in the tomographic scans. Based on these assumptions, *ε* ≈ 1.5 %. We hence predict any filament length changes across mandibular opening angles to be negligible. This prediction is supported by the observation that filament length, extracted from the tomographic scans, is independent of opening angle [LMM with random intercepts and opening angle as fixed effect: 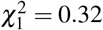, p = 0.58].

Second, we assumed that the head capsule is rigid, so that the expansion of the muscle perpendicular to the apodeme main axis remains constant. In support of this assumption, head width approximated as twice the distance of the eye to the sagittal plane was independent of opening angle [LMM with random intercepts and opening angle as fixed effect: 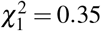, p = 0.56]. In contrast, head length decreased significantly with opening angle, but by less than 1*μ*m per degree [LMM: 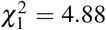, p < 0.05], amounting to a total change of only ≈ 2% across the opening range. We consider these effects small enough to be negligible.

Third, we hypothesised that the apodeme displaces only along its main axis, and that the lateral displacement, imposed by the rotation of the mandible, is bridged by the apodeme ligament. This hypothesis is supported by two observations. First, the apodeme indeed displaces only slightly along the lateral axis; the lateral projection distances of the apodeme centres-of-masses to the axis of apodeme motion never exceed 20 % of the expected lateral displacement from mandible rotation. Second, the lateral expansion of the ligament changes significantly with the sine of the apodeme angle [LMM with random intercepts and sin*γ* as fixed effect: 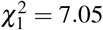, p < 0.01], accounting for around two-thirds of the expected lateral displacement. The missing third may be attributed to the biological variation between samples. We also hypothesised that any displacement of the apodeme outside the plane of rotation would be enabled by the ligament. Since the apodeme main axis and the rotational axis are basically perpendicular to each other, 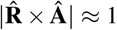, the associated out-of-plane displacement is indeed small.

### Bite force extraction for 2D sensors

Bite forces are typically measured with 1D or 2D sensors. The force components that are not measured need to be inferred from geometry. For 1D sensors, the key geometric parameter to extract the magnitude of the bite force |**F**_*b*_| is the angle between bite force vector and the sensitive axis (see Eq. 8 in the main text). For 2D sensors, |**F**_*b*_| is determined by the angle *α* between the bite force vector and the insensitive axis (**F**_*m,1*_ × **F**_*m,2*_). The orientation of the bite force vector depends on the orientations of the mandible outlever **L_*o*_** and the joint rotational axis 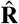, as described in the main text. The magnitude of the measured forces is equal to the components of **F**_*b*_ that are perpendicular to the insensitive axis; this relationship is characterised by the sine of *α*, i. e. the cross product between **F**_*b*_ and **F**_*m,1*_ × **F**_*m,2*_, as:

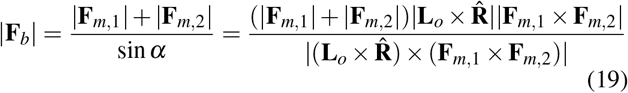

### Ant preparation for tomography

In order to characterise the morphological determinants of bite force, tomographic scans of ant heads were conducted across a maximum range of opening angles. To control mandibular opening, we built a 3D-printed device to restrain the ants after collection (see Fig. 5C). PLA rods of varying diameter were placed asymmetrically between the mandibles of the clampedant to achieve different opening angles between left and right head hemisphere. The rods were inserted into a cylindrical cavity in the device to ensure they remain in place. Subsequently, the ants were sacrificed by freezing, and their soft tissues were fixated and stained. To avoid posthumous mandible closure, the rods remained between the mandibles, but were truncated prior to scanning.

**Figure 5.**
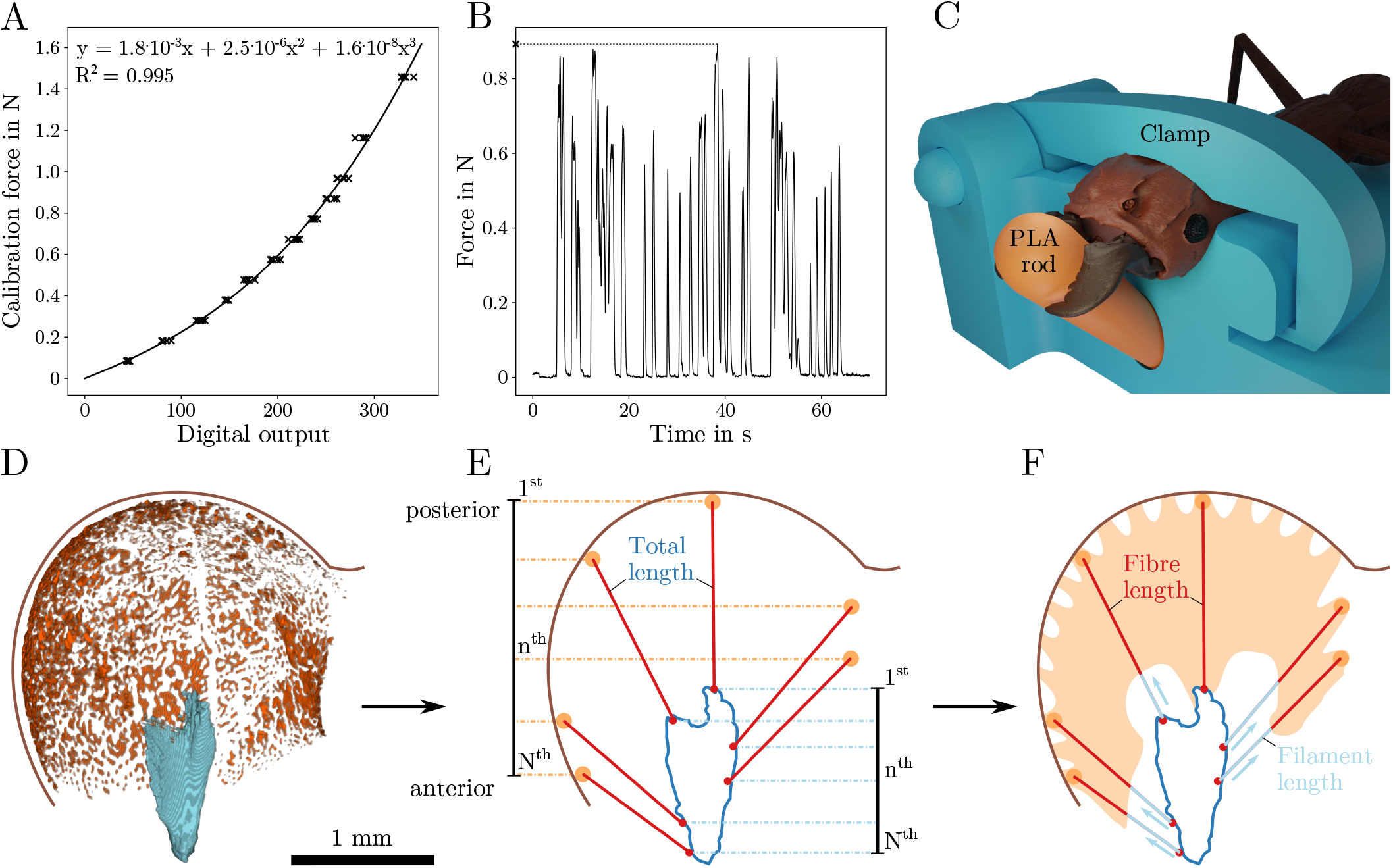
(**A**) A force sensor was calibrated to measure bite forces of leaf-cutter ants at varying mandibular opening angles. To this end, calibration weights exceeding the range of measured forces were suspended from the sensitive bite plate of the force sensor. The relationship between calibration forces and digital output was characterised using a third-degree polynomial. (**B**) Bite forces were recorded until at least five bite cycles were completed. To compare bite force across different opening angles, the maximum measured force was extracted from each force trace. (**C**) To link the measured bite forces to the morphology of the bite apparatus, we performed tomographic scans of the heads. Prior to scanning, the ants were clamped using a custom-built device; they were given PLA rods of varying thickness to bite onto. The rods prevented the mandibles from closing after death. (**D**) To extract muscle fibre length and orientation from the segmented scans, a three-step analysis was conducted. First, the fibre attachments to the head capsule were identified. (**E**) Second, according to their anterior-posterior position, these fibre attachments (N in total) were connected to the apodeme surface. (**F**) Third, from the apodeme surface, filaments were grown towards the fibre attachments until reaching muscle tissue. Fibre length was calculated as the difference between the projected total length and filament length.

The samples were imaged with an X-ray microscope using a CCD detector system with scintillator-coupled visible light optics, and tungsten transmission target. An X-ray tube voltage of 70 kV was used, and a tube current of 85 *μ*A, with an exposure of between 1000-2000 ms, and a total of 1601 projections captured over a ‘180 degrees plus fan angle’ range. An objective lens giving an optical magnification of 0.4x was selected with binning set to 2, producing isotropic voxels of 6.5 - 7.1 *μ*m. A low energy filter was placed in the beam path (LE1 – proprietary Carl Zeiss microscopy filter). The tomograms were reconstructed as 16-bit from 2D projections using a Zeiss commercial software package (XMReconstructor, Carl Zeiss), a cone-beam reconstruction algorithm based on filtered back-projection. XMReconstructor was also used to produce 2-D grey scale slices for subsequent analysis.

### Sensor calibration and bite force measurement

The sensor was calibrated by suspending calibration weights from the bite plate of the pivoting lever [see 113, for more details]. Twelve weights between 10 and 150g were used for calibration, exceeding the range of measured bite forces (109 - 1172 mN). Each weight was suspended five times, and the average sensor output over ≈ 2 s was extracted. A third degree polynomial was used to characterise the relationship between sensor output and force (*R*^2^ = 0.995, see Fig. 5A). Once calibrated, the sensor was used to measure ant bite forces (see Fig. 5B). From each force, the maximum bite force was extracted for further analysis, as described in the main text.

### Passive joint forces enabled by the opener muscle

Bite forces were measured across a large range of mandibular opening angles and the maximum stress, the shape parameter β and the optimum fibre length were extracted (see Fig 4A in the main text). The general shape of the decrease in bite force with opening angle described by the model matches well with the measurements. For large opening angles ≥ 95°, however, the model appears to underestimate the measured forces by ≈ 50 to 150mN. This difference might be attributed to passive joint forces [see 60]; elastic elements in and around the joint can act as a spring loaded by the mandible opener muscle. In order to estimate the magnitude of such joint forces, we calculated the maximum mandible opening force in analogy to Eq. 1 in the main text. The maximum stress was assumed to be the same as for the closer muscle, the physiological cross-sectional area and pennation angles were estimated using results extracted from scans of *A. vollenweideri* majors [see 81]; the effective inlever was extracted from the scans used in this study. We estimate the maximum mandibular opening force to be around 50 mN, partially bridging the mismatch between model and measurement; the opener muscle may thus indirectly contribute to an increase in bite force at large opening angles.

## References

[1] Anderson RA, McBrayer LD, Herrel A. 2008 Bite force in vertebrates: opportunities and caveats for use of a nonpareil whole-animal performance measure. Biological Journal of the Linnean Society 93: 709–720.

[2] Verwaijen D, Van Damme R, Herrel A. 2002 Relationships between head size, bite force, prey handling efficiency and diet in two sympatric lacertid lizards. Functional Ecology 16: 842–850.

[3] van der Meij M, Bout R. 2004 Scaling of jaw muscle size and maximal bite force in finches. Journal of Experimental Biology 207: 2745–2753.

[4] Gidmark NJ, Konow N, LoPresti E, Brainerd EL. 2013 Bite force is limited by the force–length relationship of skeletal muscle in black carp, *Mylopharyngodon piceus*. Biology Letters 9: 20121181.

[5] Lappin AK, Wilcox SC, Moriarty DJ, Stoeppler SA, Evans SE, Jones ME. 2017 Bite force in the horned frog (*Ceratophrys cranwelli)* with implications for extinct giant frogs. Scientific reports 7: 1–10.

[6] Herrel A, Petrochic S, Draud M. 2018 Sexual dimorphism, bite force and diet in the diamondback terrapin. Journal of Zoology 304: 217–224.

[7] Kaczmarek EB, Gidmark NJ. 2020 The bite force–gape relationship as an avenue of biomechanical adaptation to trophic niche in two salmonid fishes. Journal of Experimental Biology 223: jeb223180.

[8] Husak JF, Lappin AK, Van Den Bussche RA. 2009 The fitness advantage of a high-performance weapon. Biological Journal of the Linnean Society 96: 840–845.

[9] Hall MD, McLaren L, Brooks RC, Lailvaux SP. 2010 Interactions among performance capacities predict male combat outcomes in the field cricket. Functional Ecology 24: 159–164.

[10] Goyens J, Dirckx J, Dierick M, Van Hoorebeke L, Aerts P. 2014 Biomechanical determinants of bite force dimorphism in *Cyclommatus metallifer* stag beetles. Journal of Experimental Biology 217: 1065–1071.

[11] McLain DK, Logue J, Pratt AE, McBrayer LD. 2015 Claw-pinching force of sand fiddler crabs in relation to activity and the lunar cycle. Journal of Experimental Marine Biology and Ecology 471: 190–197.

[12] Hao W, Yao G, Zhang X, Zhang D. 2018 Kinematics and mechanics analysis of trap-jaw ant *Odontomachus monticola*. In: Journal of Physics: Conference Series. IOP Publishing, volume 986, p. 012029.

[13] Pruim G, De Jongh H, Ten Bosch J. 1980 Forces acting on the mandible during bilateral static bite at different bite force levels. Journal of Biomechanics 13: 755–763.

[14] Elner RW, Campbell A. 1981 Force, function and mechanical advantage in the chelae of the american lobster *Homarus americanus* (decapoda: Crustacea). Journal of Zoology 193: 269–286.

[15] Govind C, Blundon JA. 1985 Form and function of the asymmetric chelae in blue crabs with normal and reversed handedness. The Biological Bulletin 168: 321–331.

[16] Blundon JA. 1988 Morphology and muscle stress of chelae of temperate and tropical stone crabs *Menippe mercenaria*. Journal of Zoology 215: 663–673.

[17] Dessem D, Druzinsky RE. 1992 Jaw-muscle activity in ferrets, *Mustela putorius furo*. Journal of Morphology 213: 275–286.

[18] Levinton J, Judge M. 1993 The relationship of closing force to body size for the major claw of *Uca pugnax* (decapoda: Ocypodidae). Functional Ecology 7: 339–345.

[19] Smith LD, Palmer AR. 1994 Effects of manipulated diet on size and performance of brachyuran crab claws. Science 264: 710–712.

[20] Block JD, Rebach S. 1998 Correlates of claw strength in the rock crab, *Cancer irroratus* (decapoda, brachyura). Crustaceana 71: 468–473.

[21] Yamada SB, Boulding EG. 1998 Claw morphology, prey size selection and foraging efficiency in generalist and specialist shell-breaking crabs. Journal of Experimental Marine Biology and Ecology 220: 191–211.

[22] Herrel A, Spithoven L, Van Damme R, De Vree F. 1999 Sexual dimorphism of head size in gallotia galloti: testing the niche divergence hypothesis by functional analyses. Functional Ecology 13: 289–297.

[23] Taylor GM. 2000 Maximum force production: why are crabs so strong? Proceedings of the Royal Society of London. Series B: Biological Sciences 267: 1475–1480.

[24] Aguirre LF, Herrel A, Van Damme R, Matthysen E. 2002 Eco- morphological analysis of trophic niche partitioning in a tropical savannah bat community. Proceedings of the Royal Society of London. Series B: Biological Sciences 269: 1271–1278.

[25] Meyers JJ, Herrel A, Birch J. 2002 Scaling of morphology, bite force and feeding kinematics in an iguanian and a scleroglossan lizard. Topics in Functional and Ecological Vertebrate Morphology: 47–62.

[26] Dumont ER, Herrel A. 2003 The effects of gape angle and bite point on bite force in bats. Journal of Experimental Biology 206: 2117–2123.

[27] Erickson GM, Lappin AK, Vliet KA. 2003 The ontogeny of bite-force performance in american alligator (*Alligator mississippiensis)*. Journal of Zoology 260: 317–327.

[28] Herrel A, Podos J, Huber S, Hendry A. 2005 Evolution of bite force in darwin’s finches: a key role for head width. Journal of Evolutionary Biology 18: 669–675.

[29] Huber DR, Eason TG, Hueter RE, Motta PJ. 2005 Analysis of the bite force and mechanical design of the feeding mechanism of the durophagous horn shark *Heterodontus francisci*. Journal of Experimental Biology 208: 3553–3571.

[30] Herrel A, O’Reilly JC. 2006 Ontogenetic scaling of bite force in lizards and turtles. Physiological and Biochemical Zoology 79: 31–42.

[31] Huber DR, Weggelaar CL, Motta PJ. 2006 Scaling of bite force in the blacktip shark *Carcharhinus limbatus*. Zoology 109: 109–119.

[32] Wilson RS, Angilletta Jr MJ, James RS, Navas C, Seebacher F. 2007 Dishonest signals of strength in male slender crayfish (*Cherax dispar*) during agonistic encounters. The American Naturalist 170: 284–291.

[33] Bywater C, Angilletta Jr M, Wilson R. 2008 Weapon size is a reliable indicator of strength and social dominance in female slender crayfish *(Cherax dispar)*. Functional Ecology 22: 311–316.

[34] Claussen DL, Gerald GW, Kotcher JE, Miskell CA. 2008 Pinching forces in crayfish and fiddler crabs, and comparisons with the closing forces of other animals. Journal of Comparative Physiology B 178: 333–342.

[35] Freeman PW, Lemen CA. 2008 Measuring bite force in small mammals with a piezo-resistive sensor. Journal of Mammalogy 89: 513–517.

[36] Jones ME, Lappin AK. 2009 Bite-force performance of the last rhynchocephalian (lepidosauria: *Sphenodon*). Journal of the Royal Society of New Zealand 39: 71–83.

[37] Lailvaux SP, Reaney LT, Backwell PR. 2009 Dishonest signalling of fighting ability and multiple performance traits in the fiddler crab *Uca mjoebergi*. Functional Ecology 23: 359–366.

[38] Williams SH, Peiffer E, Ford S. 2009 Gape and bite force in the rodents *Onychomys leucogaster* and *Peromyscus maniculatus*: Does jaw-muscle anatomy predict performance? Journal of Morphology 270: 1338–1347.

[39] Pfaller J, Herrera N, Gignac P, Erickson G. 2010 Ontogenetic scaling of cranial morphology and bite-force generation in the loggerhead musk turtle. Journal of Zoology 280: 280–289.

[40] Becerra F, Echeverría A, Vassallo AI, Casinos A. 2011 Bite force and jaw biomechanics in the subterranean rodent talas tuco-tuco (*Ctenomys talarum*)(caviomorpha: Octodontoidea). Canadian Journal of Zoology 89: 334–342.

[41] Grubich JR, Huskey S, Crofts S, Orti G, Porto J. 2012 Megabites: extreme jaw forces of living and extinct piranhas (ser- rasalmidae). Scientific Reports 2: 1–9.

[42] Marshall CD, Guzman A, Narazaki T, Sato K, Kane EA, Sterba-Boatwright BD. 2012 The ontogenetic scaling of bite force and head size in loggerhead sea turtles (*Caretta caretta*): implications for durophagy in neritic, benthic habitats. Journal of Experimental Biology 215: 4166–4174.

[43] Becerra F, Casinos A, Vassallo AI. 2013 Biting performance and skull biomechanics of a chisel tooth digging rodent (*Ctenomys tuconax;* caviomorpha; octodontoidea). Journal of Experimental Zoology Part A: Ecological Genetics and Physiology 319: 74–85.

[44] Yap NW, Lin Y, Todd PA. 2013 Chelae force generation at variable gape sizes in the mud crab, *Scylla olivacea* (brachyura: Portunidae). Nature in Singapore 6: 179–185.

[45] Erickson G, Gignac P, Lappin A, Vliet K, Brueggen J, Webb G. 2014 A comparative analysis of ontogenetic bite-force scaling among crocodylia. Journal of Zoology 292: 48–55.

[46] Senawi J, Schmieder D, Siemers B, Kingston T. 2015 Beyond size - morphological predictors of bite force in a diverse insectivorous bat assemblage from malaysia. Functional Ecology 29: 1411–1420.

[47] Oka Si, Tomita T, Miyamoto K. 2016 A mighty claw: pinching force of the coconut crab, the largest terrestrial crustacean. Public Library of Science One 11: e0166108.

[48] Santana SE. 2016 Quantifying the effect of gape and morphology on bite force: biomechanical modelling and *in vivo* measurements in bats. Functional Ecology 30: 557–565.

[49] Penning DA. 2017 The scaling of bite force and constriction pressure in kingsnakes (*Lampropeltis getula*): proximate determinants and correlated performance. Integrative Zoology 12: 121–131.

[50] Malavé BM, Styga JM, Clotfelter ED. 2018 Size, shape, and sex-dependent variation in force production by crayfish chelae. Journal of Morphology 279: 312–318.

[51] Jones ME, Pistevos JC, Cooper N, Lappin AK, Georges A, Hutchinson MN, Holleley CE. 2020 Reproductive phenotype predicts adult bite-force performance in sex-reversed dragons (*Pogona vitticeps*). Journal of Experimental Zoology Part A: Ecological and Integrative Physiology 333: 252–263.

[52] South J, Madzivanzira TC, Tshali N, Measey J, Weyl OL. 2020 In a pinch: mechanisms behind potential biotic resistance toward two invasive crayfish by native african freshwater crabs. Frontiers in Ecology and Evolution 8: 72.

[53] Wheater C, Evans M. 1989 The mandibular forces ad pressures of some predacious coleoptera. Journal of Insect Physiology 35: 815–820.

[54] Paul J, Gronenberg W. 2002 Motor control of the mandible closer muscle in ants. Journal of Insect Physiology 48: 255–267.

[55] Spagna JC, Vakis AI, Schmidt CA, Patek SN, Zhang X, Tsutsui ND, Suarez AV. 2008 Phylogeny, scaling, and the generation of extreme forces in trap-jaw ants. Journal of Experimental Biology 211: 2358–2368.

[56] Heethoff M, Norton RA. 2009 A new use for synchrotron x-ray microtomography: three-dimensional biomechanical modeling of chelicerate mouthparts and calculation of theoretical bite forces. Invertebrate Biology 128: 332–339.

[57] van der Meijden A, Herrel A, Summers A. 2010 Comparison of chela size and pincer force in scorpions; getting a first grip. Journal of Zoology 280: 319–325.

[58] Huang MH. 2012 Extreme worker polymorphism in the bigheaded Pheidole ants. Ph.D. thesis, The University of Arizona.

[59] van der Meijden A, Langer F, Boistel R, Vagovic P, Heethoff M. 2012 Functional morphology and bite performance of raptorial chelicerae of camel spiders (solifugae). Journal of experimental biology 215: 3411–3418.

[60] Weihmann T, Reinhardt L, Weißing K, Siebert T, Wipfler B. 2015 Fast and powerful: Biomechanics and bite forces of the mandibles in the american cockroach *Periplaneta americana*. Public Library of Science One 10: e0141226.

[61] David S, Funken J, Potthast W, Blanke A. 2016 Musculoskeletal modelling under an evolutionary perspective: deciphering the role of single muscle regions in closely related insects. Journal of The Royal Society Interface 13.

[62] David S, Funken J, Potthast W, Blanke A. 2016 Musculoskeletal modelling of the dragonfly mandible system as an aid to understanding the role of single muscles in an evolutionary context. Journal of Experimental Biology 219: 1041–1049.

[63] Rühr PT, Edel C, Frenzel M, Blanke A. 2022 A bite force database of 654 insect species. bioRxiv.

[64] Pielström S, Roces F. 2014 Soil moisture and exvacation behaviour in the chaco leaf-cutting ant (*Atta vollenweideri*): digging performance and prevention of water inflow into the nest. Public Library of Science One 9: e95658.

[65] Bernays E. 1986 Diet-induced head allometry among foliage-chewing insects and its importance for graminivores. Science 231: 495–497.

[66] Chapman R. 1995 Mechanics of food handling by chewing insects. In: Chapman RF, de Boer G, editors, Regulatory mechanisms in insect feeding, Springer. pp. 3–31.

[67] Clissold F. 2007 The biomechanics of chewing and plant fracture: Mechanisms and implications. Advances in Insect Physiology 34.

[68] Battisti DS, Naylor RL. 2009 Historical warnings of future food insecurity with unprecedented seasonal heat. Science 323: 240–244.

[69] Dhaliwal G, Jindal V, Dhawan A. 2010 Insect pest problems and crop losses: changing trends. Indian Journal of Ecology 37: 1–7.

[70] Oliveira CM, Auad AM, Mendes SM, Frizzas MR. 2014 Crop losses and the economic impact of insect pests on brazilian agriculture. Crop protection 56: 50–54.

[71] Rosenzweig C, Elliott J, Deryng D, Ruane AC, Müller C, Arneth A, Boote KJ, Folberth C, Glotter M, Khabarov N, Neumann K, Piontek F, Pugh TAM, Schmid E, Stehfest E, Yang H, Jones JW. 2014 Assessing agricultural risks of climate change in the 21st century in a global gridded crop model intercomparison. Proceedings of the National Academy of Sciences of the U.S.A 111: 3268–3273.

[72] Deutsch CA, Tewsbury JJ, Tigchelaar M, Battisti DS, Merrill SC, Huey RB, Naylor RL. 2018 Increase in crop losses to insect pests in a warming climate. Science 361: 916–919.

[73] Crespo-Pérez V, Kazakou E, Roubik DW, Cárdenas RE. 2020 The importance of insects on land and in water: a tropical view. Current opinion in insect science 40: 31–38.

[74] Fowler HG, Pagani MI, Da Silva OA, Forti LC, Vasconelos DL. 1989 A pest is a pest is a pest? the dilemma of neotropical leafcutting ants keystone taxa of natural ecosystems. Enviromental Management 13: 671–675.

[75] Magalhães VB, Espirito Santo NB, Salles LF, Soares Jr H, Oliveira PS. 2018 Secondary seed dispersal by ants in neotropical cerrado savanna: species-specific effects on seeds and seedlings of *Siparuna guianensis* (siparunaceae). Ecological Entomology 43: 665–674.

[76] Bernays EA. 1991 Evolution of insect morphology in relation to plants. Philosophical Transactions of the Royal Society of London. Series B: Biological Sciences 333: 257–264.

[77] Bernays EA. 1998 Evolution of feeding behaviour in insect herbivores. Bioscience, JSTOR 48: 35–44.

[78] Krenn HW. 2019 Insect mouthparts: form, function, development and performance. Springer Nature.

[79] Alexander RM. 1985 The maximum forces exerted by animals. Journal of Experimental Biology 115: 231–238.

[80] Herring SW, Herring SE. 1974 The superficial masseter and gape in mammals. The American Naturalist 108: 561–576.

[81] Püffel F, Pouget A, Liu X, Zuber M, van de Kamp T, Roces F, Labonte D. 2021 Morphological determinants of bite force capacity in insects: a biomechanical analysis of polymorphic leaf-cutter ants. Journal of the Royal Society Interface 18: 20210424.

[82] Weihmann T, Kleinteich T, Gorb SN, Wipfler B. 2015 Functional morphology of the mandibular apparatus in the cockroach *Periplaneta americana* (blattodea: Blattidae) – a model species for omnivore insects. Arthropod Systematics & Phylogeny 73: 477–488.

[83] Blanke A, Machida R, Szucsich NU, Wilde F, Misof B. 2015 Mandibles with two joints evolved much earlier in the history of insects: dicondyly is a synapomorphy of bristletails, silverfish and winged insects. Systematic Entomology 40: 357–364.

[84] Blanke A. 2019 The early evolution of biting–chewing performance in hexapoda. In: Krenn HW, editor, Insect Mouthparts, Springer. pp. 175–202.

[85] Vincent JF, Wegst UG. 2004 Design and mechanical properties of insect cuticle. Arthropod structure & development 33: 187–199.

[86] Labonte D, Lenz AK, Oyen ML. 2017 On the relationship between indentation hardness and modulus, and the damage resistance of biological materials. Acta Biomaterialia 57: 373–383.

[87] Azizi E, Deslauriers A, Holt N, Eaton C. 2017 Resistance to radial expansion limits muscle strain and work. Biomechanics and modeling in mechanobiology 16: 1633–1643.

[88] Blanke A, Wipfler B, Letsch H, Koch M, Beckmann F, Beutel R, Misof B. 2012 Revival of palaeoptera—head characters support a monophyletic origin of odonata and ephemeroptera (insecta). Cladistics 28: 560–581.

[89] Gronenberg W, Brandão CRF, Dietz BH, Just S. 1998 Trap-jaws revisited: the mandible mechanism of the ant *Acanthognathus*. Physiological Entomology 23: 227–240.

[90] Gronenberg W, Hölldobler B, Alpert GD. 1998 Jaws that snap: control of mandible movements in the ant *Mystrium*. Journal of insect physiology 44: 241–253.

[91] Silva TS, Feitosa RM. 2019 Using controlled vocabularies in anatomical terminology: A case study with *Strumigenys* (hymenoptera: Formicidae). Arthropod structure & development 52: 100877.

[92] Pfuhl W. 1937 Die gefiederten muskeln, ihre form und ihre wirkungsweise. Zeitschrift für Anatomie und Entwicklungsgeschichte 106: 749–769.

[93] Gordon A, Huxley AF, Julian F. 1966 The variation in isometric tension with sarcomere length in vertebrate muscle fibres. The Journal of physiology 184: 170–192.

[94] Josephson RK. 1975 Extensive and intensive factors determining the performance of striated muscle. Journal of Experimental Zoology 194: 135–153.

[95] Williams CD, Salcedo MK, Irving TC, Regnier M, Daniel TL. 2013 The length–tension curve in muscle depends on lattice spacing. Proceedings of the Royal Society B: Biological Sciences 280: 20130697.

[96] Rockenfeller R, Günther M, Hooper SL. 2022 Muscle active force-length curve explained by an electrophysical model of interfilament spacing. Biophysical Journal 121: 1823–1855.

[97] Otten E. 1987 A myocybernetic model of the jaw system of the rat. Journal of Neuroscience Methods 21: 287–302.

[98] Wakeling JM, Tijs C, Konow N, Biewener A. 2021 Modeling muscle function using experimentally determined subject-specific muscle properties. Journal of Biomechanics 117: 110242.

[99] Otten E. 1987 Optimal design of vertebrate and insect sarcomeres. Journal of Morphology 191: 49–62.

[100] Gronenberg W, Paul J, Just S, Hölldobler B. 1997 Mandible muscle fibers in ants: fast or powerful? Cell & Tissue Research 289: 347–361.

[101] Azizi E. 2014 Locomotor function shapes the passive mechanical properties and operating lengths of muscle. Proceedings of the Royal Society B: Biological Sciences 281: 20132914.

[102] Benninghoff A, Rollhäuser H. 1952 Zur inneren mechanik des gefiederten muskels. Pflüger’s Archivfür die gesamte Physiolo- gie des Menschen und der Tiere 254: 527–548.

[103] Paul J, Gronenberg W. 1999 Optimizing force and velocity: Mandible muscle fibre attachments in ants. The Journal of Experimental Biology 202: 797–808.

[104] Paul J. 2001 Review: Mandible movements in ants. Comparative Biochemistry and Physiology Part A 131: 7–20.

[105] Sullivan S, McGechie F, Middleton K, Holliday C. 2019 3d muscle architecture of the pectoral muscles of european starling *(Sturnus vulgaris)*. Integrative Organismal Biology 1: 1–18.

[106] Rühr PT, Blanke A. 2022 forcex and forcer: a mobile setup and r package to measure and analyse a wide range of animal closing forces. Methods in Ecology and Evolution: 1–11.

[107] Cherrett JM. 1972 Some factors involved in the selection of vegetable substrate by *Atta cephalotes* (l.) (hymenoptera: Formicidae) in tropical rain forest. Journal of Animal Ecology 41: 647–660.

[108] Nichols-Orians CM, Schultz JC. 1989 Leaf toughness affects leaf harvesting by the leaf cutter ant, *Atta cephalotes* (l.) (Hy- menoptera: Formicidae). Biotropica 21: 80–83.

[109] Metscher BD. 2009 Microct for comparative morphology: simple staining methods allow high-contrast 3d imaging of diverse non-mineralized animal tissues. BMC physiology 9: 1–14.

[110] Schindelin J, Arganda-Carreras I, Frise E, Kaynig V, Longair M, Pietzsch T, Preibisch S, Rueden C, Saalfeld S, Schmid B, Tinevez J, White D, Hartenstein V, Eliceiri K, Tomancak P, Cardona A. 2012 Fiji: An open-source platform for biological-image analysis. Nature methods 9: 676–682.

[111] Yushkevich PA, Piven J, Hazlett HC, Smith RG, Ho S, Gee JC, Gerig G. 2006 User-guided 3d active contour segmentation of anatomical structures: significantly improved efficiency and reliability. NeuroImage 31: 1116–1128.

[112] Katzke J, Puchenkov P, Stark H, Economo EP. 2022 A roadmap to reconstructing muscle architecture from ct data. Integrative Organismal Biology 4: obac001.

[113] Püffel F, Roces F, Labonte D. 2022 Strong positive allometry of bite force in leaf-cutter ants increases the range of cuttable plant tissues. bioRxiv.

[114] Larabee FJ, Gibson JC, Rivera MD, Anderson PS, Suarez AV. 2022 Muscle fatigue in the latch-mediated spring actuated mandibles of trap-jaw ants. Integrative and Comparative Biology.

[115] Daly DC, Mercier PP, Bhardwaj M, Stone AL, Aldworth ZN, Daniel TL, Voldman J, Hildebrand JG, Chandrakasan AP. 2009 A pulsed uwb receiver soc for insect motion control. IEEE Journal of solid-state circuits 45: 153–166.

[116] Cao F, Zhang C, Vo Doan TT, Li Y, Sangi DH, Koh JS, Huynh NA, Aziz MFB, Choo HY, Ikeda K, et al. 2014 A biological micro actuator: graded and closed-loop control of insect leg motion by electrical stimulation of muscles. Public Library of Science One 9: e105389.

[117] Choo HY, Li Y, Cao F, Sato H. 2016 Electrical stimulation of coleopteran muscle for initiating flight. Public Library of Science One 11: e0151808.

[118] Field A, Miles J, Field Z. 2012 Discovering statistics using r. Great Britain: Sage Publications, Ltd.

[119] Richter A, Garcia FH, Keller RA, Billen J, Katzke J, Boudinot BE, Economo EP, Beutel RG. 2021 The head anatomy of *Protanilla lini* (hymenoptera: Formicidae: Leptanillinae), with a hypothesis of their mandibular movement. Myrmecological News 31: 85–114.

[120] Guschlbauer C, Scharstein H, Büschges A. 2007 The extensor tibiae muscle of the stick insect: biomechanical properties of an insect walking leg muscle. Journal of Experimental Biology 210: 1092–1108.

[121] Brainerd EL, Azizi E. 2005 Muscle fiber angle, segment bulging and architectural gear ratio in segmented musculature. Journal of Experimental Biology 208: 3249–3261.

[122] Azizi E, Brainerd EL, Roberts TJ. 2008 Variable gearing in pennate muscles. Proceedings of the National Academy of Sciences 105: 1745–1750.

[123] Lappin AK, Jones ME. 2014 Reliable quantification of bite-force performance requires use of appropriate biting substrate and standardization of bite out-lever. Journal of Experimental Biology 217: 4303–4312.

[124] Full R, Ahn A. 1995 Static forces and moments generated in the insect leg: comparison of a three-dimensional musculo-skeletal computer model with experimental measurements. The journal of Experimental Biology 198: 1285–1298.

[125] Barbosa P, Berry D, Kary CK. 2015 Insect histology: Practical laboratory techniques. John Wiley & Sons.

[126] Zachar J, Zacharová D. 1966 The length-tension diagram of single muscle fibres of the crayfish. Experientia 22: 451–452.

[127] April EW, Brandt PW. 1973 The myofilament lattice: Studies on isolated fibers: Iii. the effect of myofilament spacing upon tension. The Journal of General Physiology 61: 490–508.

[128] Chapple WD. 1983 Mechanical responses of a crustacean slow muscle. Journal of Experimental Biology 107: 367–383.

[129] Ahn AN, Meijer K, Full RJ. 2006 In situ muscle power differs without varying in vitro mechanical properties in two insect leg muscles innervated by the same motor neuron. Journal of Experimental Biology 209: 3370–3382.

[130] Josephson RK, Stokes DR. 1987 The contractile properties of a crab respiratory muscle. Journal of Experimental Biology 131: 265–287.

[131] Tu MS, Daniel TL. 2004 Submaximal power output from the dorsolongitudinal flight muscles of the hawkmoth *Manduca sexta*. Journal of Experimental Biology 207: 4651–4662.

[132] Shimomura T, Iwamoto H, Doan TTV, Ishiwata S, Sato H, Suzuki M. 2016 A beetle flight muscle displays leg muscle microstructure. Biophysical Journal 111: 1295–1303.

[133] Onoda Y, Westoby M, Adler PB, Choong AMF, Clissold FJ, Cornelissen JHC, Díaz S, Dominy NJ, Elgart A, Enrico L, Fine PVA, Howard JJ, Jalili A, Kitajima K, Kurokawa H, McArthur C, Lucas PW, Markesteijn L, Pérez-Harguindeguy N, Poorter L, Richards L, Santiago LS, Sosinski Jr EE, Van Bael SA, Warton DI, Wright IJ, Joseph Wright S, Yamashita N. 2011 Global patterns of leaf mechanical properties. Ecology Letters 14: 301–312.

[134] Tautz J, Roces F, Hölldobler B. 1995 Use of a sound-based vibratome by leaf-cutting ants. Science 267: 84.

[135] Schofield RM, Emmett KD, Niedbala JC, Nesson M. 2011 Leafcutter ants with worn mandibles cut half as fast, spend twice the energy, and tend to carry instead of cut. Behavioral Ecology and Sociobiology 65: 969–982.

[136] Freeman PW, Lemen CA. 2010 Simple predictors of bite force in bats: the good, the better and the better still. Journal of Zoology 282: 284–290.

[137] Habegger ML, Motta PJ, Huber DR, Dean MN. 2012 Feeding biomechanics and theoretical calculations of bite force in bull sharks (*Carcharhinus leucas*) during ontogeny. Zoology 115: 354–364.

[138] Lowie A, De Kegel B, Wilkinson M, Measey J, O’Reilly JC, Kley NJ, Gaucher P, Brecko J, Kleinteich T, Adriaens D, et al. 2022 The relationship between head shape, head musculature and bite force in caecilians (amphibia: Gymnophiona). Journal of Experimental Biology 225: jeb243599.

[139] Erickson GM, Kirk SDV, Su J, Levenston ME, Caler WE, Carter DR. 1996 Bite-force estimation for tyrannosaurus rex from tooth-marked bones. Nature 382: 706–708.

[140] Santana SE, Dumont ER, Davis JL. 2010 Mechanics of bite force production and its relationship to diet in bats. Functional Ecology 24: 776–784.

[141] Sakamoto M, Ruta M, Venditti C. 2019 Extreme and rapid bursts of functional adaptations shape bite force in amniotes. Proceedings of the Royal Society B 286: 20181932.

[142] Li D, Zhang K, Zhu P, Wu Z, Zhou H. 2011 3d configuration of mandibles and controlling muscles in rove beetles based on micro-ct technique. Analytical and bioanalytical chemistry 401: 817–825.

[143] Lillico-Ouachour A, Metscher B, Kaji T, Abouheif E. 2018 Internal head morphology of minor workers and soldiers in the hyperdiverse ant genus *Pheidole*. Canadian Journal of Zoology 96: 383–392.

[144] Richter A, Keller RA, Rosumek FB, Economo EP, Garcia FH, Beutel RG. 2019 The cephalic anatomy of workers of the ant species *Wasmannia affinis* (formicidae, hymenoptera, insecta) and its evolutionary implications. Arthropod structure & development 49: 26–49.

[145] Richter A, Garcia FH, Keller RA, Billen J, Economo EP, Beutel RG. 2020 Comparative analysis of worker head anatomy of *Formica* and *Brachyponera* (hymenoptera: Formicidae). Arthropod Systematics & Phylogeny 78: 133–170.

[146] Zhang W, Li M, Zheng G, Guan Z, Wu J, Wu Z. 2020 Multifunctional mandibles of ants: Variation in gripping behavior facilitated by specific microstructures and kinematics. Journal of Insect Physiology 120: 103993.

[147] van de Kamp T, Mikó I, Staniczek AH, Eggs B, Bajerlein D, Faragó T, Hagelstein L, Hamann E, Spiecker R, Baumbach T, et al. 2022 Evolution of flexible biting in hyperdiverse parasitoid wasps. Proceedings of the Royal Society B 289: 20212086.

